# Adeno-Associated Viral Tools to Trace Neural Development and Connectivity Across Amphibians

**DOI:** 10.1101/2024.02.15.580289

**Authors:** Eliza C.B. Jaeger, David Vijatovic, Astrid Deryckere, Nikol Zorin, Akemi L. Nguyen, Georgiy Ivanian, Jamie Woych, Rebecca C. Arnold, Alonso Ortega Gurrola, Arik Shvartsman, Francesca Barbieri, Florina A. Toma, Gary J. Gorbsky, Marko E. Horb, Hollis T. Cline, Timothy F. Shay, Darcy B. Kelley, Ayako Yamaguchi, Mark Shein-Idelson, Maria Antonietta Tosches, Lora B. Sweeney

## Abstract

The development, evolution, and function of the vertebrate central nervous system (CNS) can be best studied using diverse model organisms. Amphibians, with their unique phylogenetic position at the transition between aquatic and terrestrial lifestyles, are valuable for understanding the origin and evolution of the tetrapod brain and spinal cord. Their metamorphic developmental transitions and unique regenerative abilities also facilitate the discovery of mechanisms for neural circuit remodeling and replacement. The genetic toolkit for amphibians, however, remains limited, with only a few species having sequenced genomes and a small number of transgenic lines available. In mammals, recombinant adeno-associated viral vectors (AAVs) have become a powerful alternative to genome modification for visualizing and perturbing the nervous system. AAVs are DNA viruses that enable neuronal transduction in both developing and adult animals with low toxicity and spatial, temporal, and cell-type specificity. However, AAVs have never been shown to transduce amphibian cells efficiently. To bridge this gap, we established a simple, scalable, and robust strategy to screen AAV serotypes in three distantly-related amphibian species: the frogs *Xenopus laevis* and *Pelophylax bedriagae,* and the salamander *Pleurodeles waltl,* in both developing larval tadpoles and post-metamorphic animals. For each species, we successfully identified at least two AAV serotypes capable of infecting the CNS; however, no pan-amphibian serotype was identified, indicating rapid evolution of AAV tropism. In addition, we developed an AAV-based strategy that targets isochronic cohorts of developing neurons – a critical tool for parsing neural circuit assembly. Finally, to enable visualization and manipulation of neural circuits, we identified AAV variants for retrograde tracing of neuronal projections in adult animals. Our findings expand the toolkit for amphibians to include AAVs, establish a generalizable workflow for AAV screening in non-canonical research organisms, generate testable hypotheses for the evolution of AAV tropism, and lay the foundation for modern cross-species comparisons of vertebrate CNS development, function, and evolution.

## Introduction

The study of the amphibian nervous system has enhanced our knowledge of the development and evolution of neural cell types. Historically, amphibians were instrumental in the discovery of the Spemann-Mangold embryonic organizer (reviewed in ^1^) and of fundamental mechanisms of synaptic transmission ^2^ More recent work on neuronal cell type determination, sensory processing, motor pattern generation, circuit formation, regeneration and plasticity ^3–13^, as well as human neurodevelopmental disorders ^14,15^ has benefited greatly from the use of amphibian models. Finally, the amphibian’s position in vertebrate evolution also enables inferences about nervous system adaptations across the evolutionary transition from aquatic to terrestrial environments ^8,16–18^.

Amphibians offer several experimental advantages. Development is external, clutch sizes are large, species can be obtained from stock centers ^19–21^ or commercial suppliers, adults may be bred in the laboratory or studied in the wild ^22,23^. These features have facilitated recent expansion of the genetic toolkit in frogs and salamanders, including transgenesis and CRISPR/Cas-based genome editing ^24–28^. However, the long generation time of frogs and salamanders, reaching sexual maturity after at least one year post-fertilization ^22,29,30^, limits the application of standard approaches used in more established genetic model organisms such as worms, flies, fish, and mice.

To move from descriptive to functional studies in amphibians, we require a toolkit to manipulate gene expression rapidly with spatiotemporal control. Electroporation can be used for the acute delivery of genes into amphibian cells, but it suffers from low efficacy and transient expression, with cell-type specificity additionally limited by the availability of cis-regulatory elements ^31,32^. Direct injection of RNAs or morpholinos into developing frogs briefly perturbs gene expression, but its effect is also transient and only effective at early developmental stages ^14,33^. Viral vectors in contrast would offer an efficient and stable alternative that could be tailored to target cell types specifically ^34–36^.

As part of a larger effort to understand how diverse cell types assemble and function within the vertebrate central nervous system (CNS), a variety of viral vectors have been developed over the last decades. Many that were first identified to study the nervous system, including the rabies and pseudorabies viruses, had limited utility due to their high toxicity ^37^. Other viruses that have been employed to express transgenes in mammalian neural circuits include herpes simplex virus, vesicular stomatitis virus (VSV), lentivirus, sindbis virus ^38^, adenovirus, and adeno-associated virus (AAV) ^39^.

However, despite their demonstrated utility in mammals, viral vectors have been difficult to implement in amphibians. In frogs, transduction of neurons with vaccinia virus, adenoviruses, recombinant rabies virus, and VSV is possible but limited by either cytotoxicity, high biosafety risk, or availability ^40–43^. Transduction with recombinant AAVs or lentiviruses in *Xenopus* frogs has thus far been unreliable or entirely unsuccessful ^42^. In axolotl, foamy viruses, pseudotyped baculoviruses, vaccinia viruses and retroviruses have been shown to infect intact or regenerating tissue ^44–48^, but either they have a potent effect on cell health, their expression is restricted to proliferating cells or they are not commercially available, limiting utility. In other amphibian species, viral tools are largely untested and present a significant opportunity for expanding the toolkit.

AAVs, a member of the Parvoviridae family, are single-stranded DNA dependoviruses that can be rendered completely replication-defective recombinantly. AAVs are widely used in neuroscience and gene therapy research, due to their ease of generation, low immunogenicity, and minimal biosafety risk level ^49–52^. These viral vectors exist in different natural and engineered capsid serotypes, with some tissue and cell-type selectivity ^53^. Further cell-type and spatial restriction of AAV expression can be achieved with promoters or enhancers, intersectional strategies (e.g. Cre-lox or Flp-FRT recombination) ^49,54^, or by targeting a specific developmental stage ^55,56^. In addition, AAV-driven gene expression is stable and lasts for long periods of time without causing a substantial decline in neuronal health ^57^. Due to these advantages, the number and availability of AAVs has expanded dramatically, allowing for the visualization and manipulation of neural circuits via antero- and retrograde tracing, calcium indicators, and actuators, without the need for transgenic lines.

Given the advantages of AAVs over other viral vectors, we chose to screen commercially available AAV serotypes in two frog species: the African clawed frog *Xenopus laevis* and the Levant water frog *Pelophylax bedriagae*, and in a salamander species, the Iberian ribbed newt *Pleurodeles waltl*. *Xenopus* is an established model for the study of neural circuit development and function. *Pelophylax,* an Eastern Mediterranean frog, is an emerging model in the fields of population genetics and ecology, abundant in the field but not yet captive-bred. *Pleurodeles* is popular in regeneration research, and is an emerging model for studies on neural circuit development, function, and evolution. These three species are representatives of distantly-related branches of the amphibian tree (**Figure 1A**). Frogs (order: Anura) and salamanders (order: Caudata) are, together with Cecilians, the three orders of modern amphibians, and diverged about 272 million years ago ^58,59^. *Xenopus* and *Pelophylax* frogs belong to two distantly-related families - Pipidae and Ranidae, respectively - which diverged in the Early Jurassic, about 182.5 million years ago ^58,59^. The phylogenetic position of these three species allows us to test whether conserved principles of AAV tropism exist between distantly-related amphibians.

**Figure 1.**
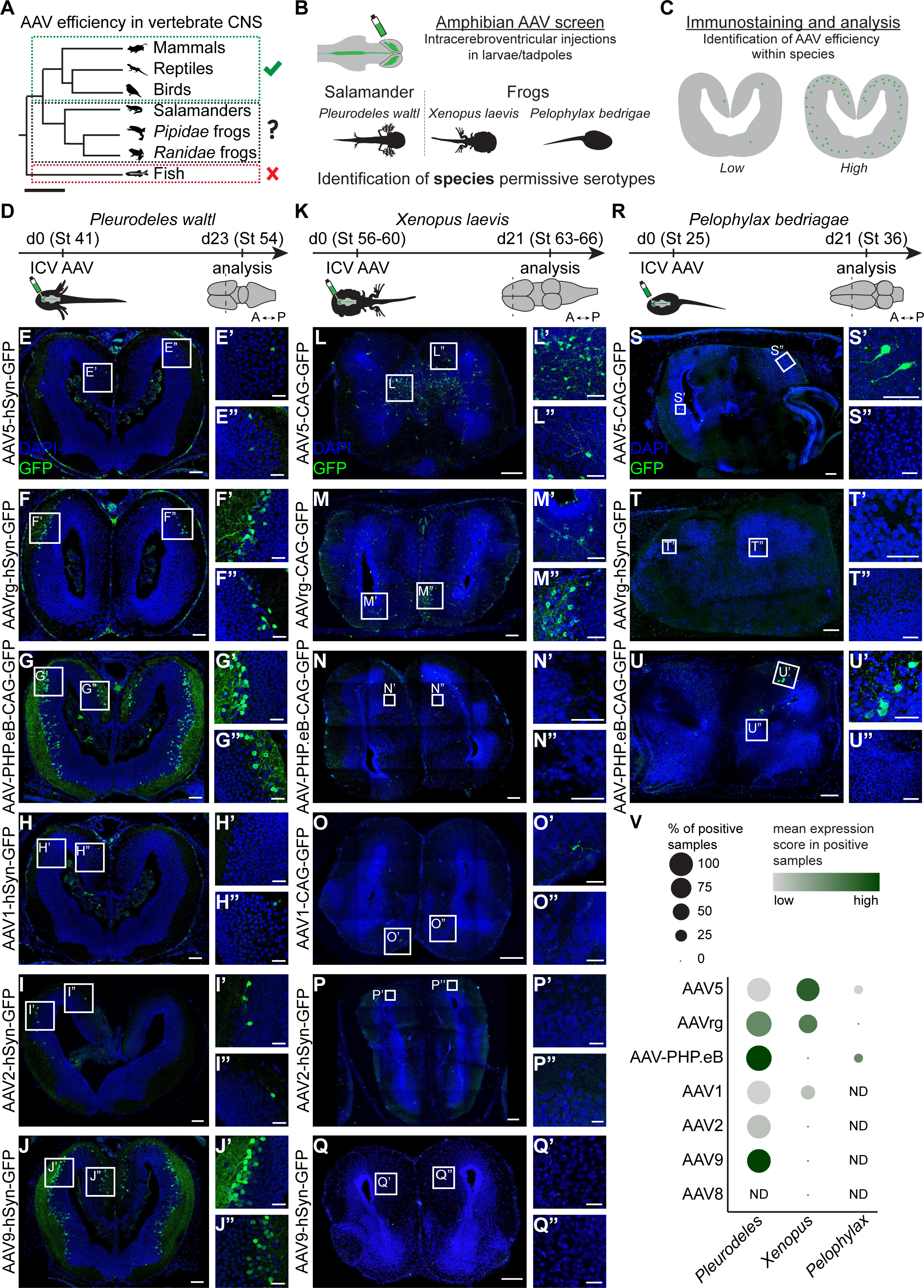
AAV serotype screening in *Pleurodeles*, *Xenopus* and *Pelophylax* larvae. **A.** Overview of AAV use across vertebrate model organisms. Check mark indicates successful transduction of any AAV serotype in the CNS, whereas the “x” indicates that no tested serotype so far has been able to infect the CNS. Question mark indicates the need for further investigation. Scale bar represents 200 MYA (million years ago). **B.** *Pleurodeles, Xenopus* and *Pelophylax* larvae were injected with a panel of AAV serotypes in the cerebral ventricle, allowing for the identification of species-specific AAV expression patterns. **C.** Three weeks after injection, brain samples were sliced and GFP signal was amplified. Transduction efficiency was scored from absent over low to high (see **STAR Methods**). **D-U.** Transduction efficiency of different AAV serotypes was analyzed after intracerebroventricular injection in *Pleurodeles waltl* (**D-J**), *Xenopus laevis* (**K-Q**) and *Pelophylax bedriagae* (**R-U**) larvae, at the developmental stages indicated in **D**, **K**, and **R**. For each species, panels on the left show representative overview images of coronal sections through the telencephalon, and boxes indicate magnified regions on the right. Scale bars in overview images and magnifications are 100 μm and 40 μm, respectively. AAV-driven GFP, green. DAPI, blue. **V.** Dot plot representing the transduction efficiency of each serotype, scored independently in each species (see **STAR Methods** for scoring criteria). Dot size represents the percentage of animals in which the injection resulted in labeling of any number of cells, and color reflects the average efficiency of viral labeling in positive samples only. **Abbreviations:** A, anterior; CNS, central nervous system; d, day; ICV, intracerebroventricular; ND, not determined; P, posterior; St, stage.

Here, we establish a rapid AAV screening strategy in amphibians and identify serotypes that successfully infect neurons in each of the three species. Furthermore, we introduce novel neuroscience applications of AAVs in developing and mature amphibians. First, we show that AAVs injected in larvae reproducibly transduce isochronic cohorts of neurons. Second, we find that AAV injections in post-metamorphic amphibians label neurons and their axonal projections. These results open up new opportunities for labeling and manipulating both developing and mature neural circuits in amphibians and define a generalizable roadmap for establishing AAV tools in new vertebrate species.

## Results

### An efficient AAV screening strategy in amphibians

To examine whether existing AAV serotypes can drive expression of transgenes in the amphibian CNS *in vivo*, we sought to establish a simple, scalable, and robust screening strategy across all three species of interest (**Figure 1A-C**). Inspired by the widespread AAV uptake in the developing mouse brain ^56,60,61^, we focused on larvae and tadpoles, phases of the amphibian life cycle where simple functional larval neural circuits coexist with neural progenitors and developing neurons of adult circuits ^62^.

For a simple, scalable, and robust screen, we chose intracerebroventricular injections of AAVs in larvae. This injection technique is simple and minimally invasive, because the larval CNS ventricles are visible (larvae are translucent) ^63^ and accessible through the skin and cartilage. Additionally, using larvae made our screen easy to scale to multiple serotypes, because larvae are abundant and easy to maintain. Furthermore, AAVs injected into the ventricular system reach a large number of brain regions and neurons, increasing the probability of detecting labeled cells for serotypes with very low transduction efficiency. To make our screen more robust, we compared AAV serotypes driving the expression of the same fluorophore (the enhanced green fluorescent protein GFP), and amplified the GFP signal by immunohistochemistry. Finally, we selected a broad panel of AAV serotypes - AAV1, AAV2, AAV5, AAV8, AAV9, AAVrg or AAV-PHP.eB - with distantly-related capsid protein sequences ^64^, to ensure broad coverage of AAV diversity.

### AAVs transduce the developing brain in salamanders

In the salamander *Pleurodeles waltl*, we targeted our screen to early active larvae (stage 41) ^65^. We injected AAV1, AAV2, AAV5, AAV9, AAVrg or AAV-PHP.eB into the telencephalic ventricle of one hemisphere and observed rapid spread of the injected solution throughout the ventricle. To ensure neuronal-specific labeling, all AAV vectors tested in *Pleurodeles* - except AAV-PHP.eB - drove GFP expression under control of the pan-neuronal human synapsin 1 (hSyn) promoter ^66^. For the AAV-PHP.eB serotype, a vector carrying GFP under the ubiquitous CAG promoter - a synthetic promoter including the cytomegalovirus early enhancer element and chicken beta-actin promoter ^67^ - was used.

Three weeks after injection, we evaluated the transduction efficiency of all serotypes in the forebrain and midbrain (**Figure 1D-J** and **Figure S1A-I**). We observed on average a low number of GFP positive cells (0-5 cells per brain section) after AAV1, AAV2 and AAV5 infection, a moderate number (6-25 cells per section) after AAVrg infection, and a high number of GFP positive cells (>25 cells per section) after AAV9 and AAV-PHP.eB infection. Labeling was reproducible across individuals, except in AAV2-injected larvae, where we observed higher animal-to-animal variability in transduction efficiency (**Table S1**). GFP+ cells generally occupied a distinct layer of cells in the mantle zone, which primarily includes neurons. Labeling spanned both the forebrain and midbrain, consistent with the spread of the viral suspension through the ventricles and transduction of cells across the brain (**Figure S1B-I**). The morphology of labeled cells was consistent with their neuronal nature: GFP labeled not only cell bodies but also dendrites, axonal tracts, and brain commissures (**Figure 1G,J**). These results indicate that many serotypes transduce neurons in developing salamanders, and AAV9, AAVrg, and AAV-PHP.eB do so with high efficiency.

### AAVs transduce the developing CNS of Xenopus laevis frogs

We next evaluated the transduction efficiency of AAV1, AAV2, AAV5, AAV8, AAV9, AAVrg, and AAV-PHP.eB in the frog *Xenopus laevis*. Prometamorphic tadpoles (NF stage 56-60) ^68^ were chosen for the primary screen since the brain organization and immune system at this stage are similar to juvenile and adult frogs ^69–71^. Prior to NF stage 60, *Xenopus* tadpole skulls are largely cartilaginous ^72^, allowing easy access to the brain ventricle through the dorsal cranium. The virus administered via the midbrain ventricle spread into other ventricles, maximizing the number of brain areas screened for AAV transduction.

Three weeks after intracerebroventricular injections, we observed no GFP labeling with AAV2, AAV8, AAV9 and AAV-PHP.eB injections; a low number of GFP positive cells (0-5 cells per section) with AAV1 injections; and a high number of GFP positive cells (>25 cells per section) with AAV5 and AAVrg injections (**Figure 1K-Q, Table S1**). AAV5 and AAVrg-transduced cells were widespread throughout the telencephalon (**Figure 1K-Q**); mesencephalon (**Figure 2**); diencephalon, and rostral spinal cord **(Figure S2)**.

**Figure 2.**
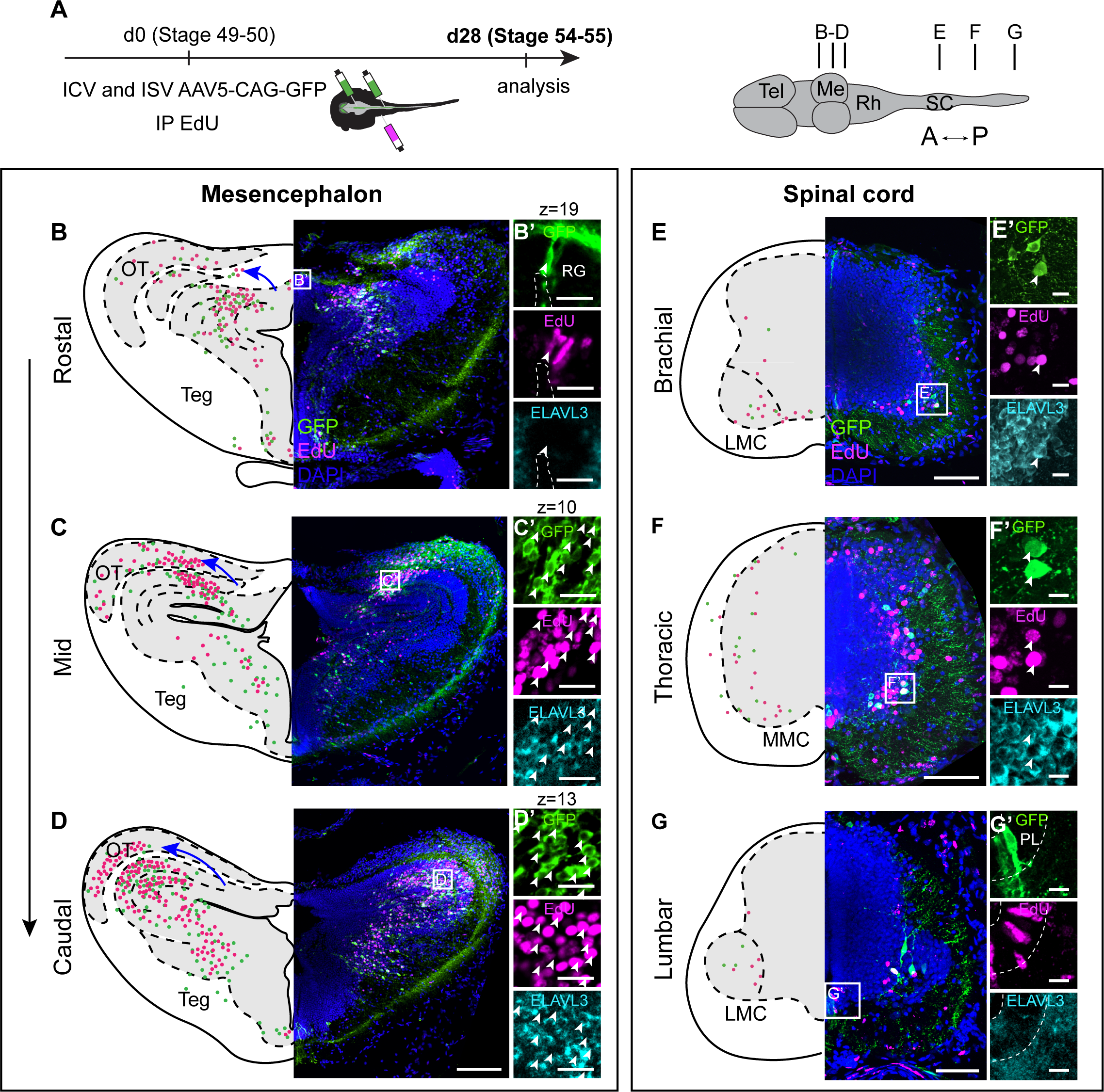
AAV labels distinct isochronic cohorts of neurons across the frog CNS. **A.** Experimental design, illustrating intracerebro- and intraspinoventricular injections of AAV5-CAG-GFP, coupled with intraperitoneal injections of EdU in prometamorphic *Xenopus laevis* tadpoles. The tissue was analyzed three weeks later, and the GFP and EdU signals were visualized. Representative coronal (brain) or transverse (spinal cord) sections (40 µm thick) are shown for different regions of the *Xenopus* CNS. Section levels are indicated on the schematic on the right. **B-D.** AAV and EdU signals overlap at the regional level at different rostral-caudal levels of the mesencephalon, marking the proliferating wedges of tectal neurons. Expansion of frog tectum starts in the medial zone at the rostral end and progresses laterally in more caudal regions as indicated by the blue arrow. Insets show co-localization of EdU, AAV and the neuronal marker Elavl3 in transduced neurons (**C’-D’**) and labeling of the radial glia with AAV and EdU but not Elavl3 (**B’**). **E-G.** In the spinal cord, AAV labeling captures the expansion of either LMC motor neurons at limb levels (**E,G**) or ventral and dorsal interneuron and MMC motor neurons at the thoracic level (**F**). AAV transduces both neurons (**E’, F’**) and radial glia (**G’**), many of which are also EdU-positive. For brain sections, scale bars in overview images and magnifications are 400 µm and 20 µm in **B-D**, and 100 µm and 20 µm in **E-G**. Images show maximum intensity projections of the entire 40 µm thick z-stack except in **B’**,**C’**, and **D’** where a single z-plane is shown due to the large amounts of labeling. AAV-driven GFP, green. EdU, magenta. Elavl3/4, cyan. DAPI, blue. **Abbreviations:** A, anterior; d, day; ICV, intracerebroventricular injection; IP, intraperitoneal injection; ISV, intraspinoventricular injection; LMC, lateral motor column; Me, mesencephalon; MMC, medial motor column; OT, optic tectum; P, posterior; PL, progenitor layer; RG, radial glia; Rh, rhombencephalon; Sc, spinal cord; Teg, tegmentum; Tel, telencephalon.

Both neuronal hSyn and ubiquitous CAG promoters were tested in *Xenopus* to assess the robustness and neuronal tropism of each promoter in driving GFP expression. For both AAV5 and AAVrg, expression driven by the hSyn promoter labeled similar numbers of cells as the CAG promoter. However, the synthetic CAG promoter drove higher expression levels than the hSyn promoter, with more brightly labeled soma and axons (**Figure S1K** versus **S1L**). Co-staining with Elavl3/4 (HuC/D), which selectively marks most neurons in the CNS ^73^, was used to evaluate whether the labeled cells were neurons (**Figure S1J-L**). AAV5 and AAVrg with the hSyn and CAG promoters both induced efficient neuronal labeling, with an average of ∼65% of AAV+ cells expressing the neuronal marker across conditions **(Figure S1M**).

We then expanded our screening to an earlier stage: the larval NF41 stage, characterized by escape swim behavior and a smaller and simpler developing CNS ^74^. Intracerebroventricular injection at NF41 demonstrated that the same serotypes - AAV1, AAV5, and AAVrg - that successfully transduce neurons at NF stage 56-60 also do so at earlier embryonic stages (**Figure S1N-Q**). This supports the conclusion that AAV transduces neurons across *Xenopus* tadpole development, independent of the maturity of the immune and nervous systems.

In a separate set of experiments in *Xenopus* tadpoles, we screened AAVs in the spinal cord. Hypothesizing that the AAV serotypes that efficiently transduce the brain would also transduce the spinal cord, we injected AAV5-CAG-GFP and AAVrg-CAG-GFP into the central canal (intraspinoventricular) at NF stage 57-60 (**Figure S2E-H**). As in the brain, we found that AAV5 and AAVrg variants sparsely transduced neurons at all levels of the spinal cord, demonstrating the ability of these serotypes to label neurons across the CNS.

Finally, to assess whether AAVs transduce other frog cell types besides neurons, we tested an immortalized *X. laevis* cell line, XLJ-1. Using the XLJ-1 line, we evaluated the transduction efficiency of AAVs over time *in vitro*. Out of six AAV vectors screened, AAV1-CAG-GFP efficiently transduced XLJ-1 cells after five days (**Figure S1R**). This result indicates that AAVs have potential to target other cell types and tissues in amphibians.

### AAVs transduce the developing CNS of Pelophylax bedriagae frogs

To test the efficiency of AAV serotype transduction across distantly-related frogs, we screened AAVs in wild-caught Levant water frogs, *Pelophylax bedriagae*, via intracerebroventricular injections at developmental larval stages 24-27 ^75,76^. After AAV5 and AAV-PHP.eB injection, we observed sparse but clear GFP expression in the telencephalon, with brightly marked neuronal soma and dendrites (**Figure 1R,S,U**). AAV5 injection led to somewhat more limited reporter gene expression than AAV-PHP.eB injection (1 to 5 cells per section, and 6 to 10 cells per section, respectively; **Figure 1S,U**). No reporter gene expression was detected for AAVrg (**Figure 1T**). In contrast to what was observed in *Xenopus* and *Pleurodeles*, the experiments in *Pelophylax* resulted in higher inter-individual variability (**Figure 1V**). These results indicate that, while expression in *Pelophylax* tadpoles was sparse and variable, AAVs can successfully infect telencephalic neurons across Anurans, even from wild-caught animals.

In summary, comparing across *Pleurodeles*, *Xenopus* and *Pelophylax* amphibian species, we identified at least two serotypes per species able to transduce neurons in the developing brain. Although these three species were all susceptible to AAV infection, we observed differences in the efficiency of transduction (average number of cells transduced per brain) as well as in the inter-individual variability (**Figure 1V**). In *Pleurodeles*, the best performing serotypes were AAVrg, AAV-PHP.eB, and AAV9; in *Xenopus*, AAV5 and AAVrg; and in *Pelophylax*, AAV5 and AAV-PHP.eB. AAVrg was the only serotype able to transduce both *Xenopus* and *Pleurodeles* neurons, and none of the serotypes tested produced reporter expression in all three amphibian species.

### AAVs target cohorts of developing neurons with temporal specificity

Intracerebroventricular administration of AAVs in both *Xenopus* and *Pleurodeles* larvae resulted in reproducible labeling of specific cell clusters in each brain region (**Figure 1**). This observation motivated us to probe further the rules and patterns of AAV transduction during larval development. In mouse embryos, intraventricular AAV injection labels isochronic cohorts of entorhinal cortex and hippocampal neurons, defined as groups of neurons that develop at the same time, with synchronous maturation and assembly into distinct neural circuits ^56^. We hypothesized that AAV may target isochronic cohorts of neurons in developing amphibians as well, explaining the reproducible patterns of GFP expression obtained after intracerebroventricular injections. The ability to target distinct isochronic cohorts of neurons using AAVs would allow for analyses of the relationships between neuronal birth order, circuit assembly, and function throughout the developing CNS. This would provide a critical tool to target, visualize and manipulate specific neuronal subpopulations during CNS development, remodeling, and regeneration.

### AAVs label isochronic cohorts of developing Xenopus laevis neurons

Previous birthdating studies in *Xenopus laevis* mapped precisely when during development each brain region is generated ^62,77,78^, an ideal starting point to test whether AAVs are taken up specifically by developing neurons or their progenitors. To address this question, we employed simultaneous intracerebroventricular or intraspinoventricular injection of AAV5-CAG-GFP, and intraperitoneal injection of EdU, a thymidine analog that incorporates into cells in the S-phase of the cell cycle ^79,80^, during premetamorphic NF49-50 tadpole stages (**Figure 2A**). Three weeks following dual AAV-EdU injection, we evaluated the number and distribution of EdU+, GFP+, and EdU+/GFP+ cells in the progenitor and mantle zones of the brain and spinal cord.

EdU labeling was pronounced in the optic tectum (**Figure 2B-D**) and spinal cord (**Figure 2E-G**), as well as the forebrain and thalamus (**Figure S2C-D**), consistent with previous studies ^78,81–85^. The spatial distribution of GFP+ cells matched the distribution of EdU+ cells (**Figure 2B-G** and **Figure S2C-D**). Across the CNS, ∼25% of transduced cells on average were also positive for EdU. Regionally, of all transduced cells, ∼20% were EdU positive in the telencephalon, ∼25% in the diencephalon, ∼40% in mesencephalon, and ∼20% in the spinal cord (**Figure S2B**). Other AAV-transduced cells, although not co-labeled by EdU, were intermingled with EdU+ cells. Conversely, regions that were largely EdU negative, and thus minimally proliferating at the time of injection, such as the hypothalamus as compared to adjacent thalamus (**Figure S2D**), were GFP negative.

In the optic tectum, the EdU labeling pattern closely followed the established course of tectum development during these stages of metamorphosis ^85^. Expansion of the optic tectum involves the serial addition of strips or “wedges” of neurons, medially forming segments that extend from the ventricular to the pial surface ^84,85^. The AAV labeling pattern recapitulated this zoned proliferation. Within the tectum, we observed that AAV and EdU labeling clustered within the same mediolateral and dorsoventral zones (**Figure 2B-D**), suggesting both populations of neurons were born around the time of injection.

During frog metamorphosis, the spinal cord also undergoes extensive transformation with timed waves of neurogenesis yielding new molecularly and spatially distinct cell types ^62^. To assess whether AAVs could be used to access isochronic cohorts in the spinal cord, we injected AAV5-CAG-GFP intraspinoventricularly at early (NF49-50), mid (NF52) or late (NF57) metamorphic stages. Early metamorphic injections preferentially transduced the ventral spinal cord (**Figure 2E-G**), intermediate stage injections labeled both ventral and dorsal populations (**Figure S2E-F**), while late injections labeled the dorsal spinal cord (**Figure S2G-H**).

Next, we sought to evaluate whether the AAV labeling of ventral versus dorsal spinal cord at different time points correlated with a difference in cell proliferation by paired AAV5-CAG-GFP and EdU injections (**Figure 2A)**. At the early NF49-50 stage of metamorphosis, we found that GFP and EdU both localized to the same spinal region (**Figure 2E-G)**. At brachial/lumbar limb-level segments, the majority of GFP+, EdU+, or co-labeled neurons were situated in the ventral horn, including lateral motor neurons identified by their distinctive settling position and morphology (**Figure 2E,G**). At thoracic torso-level segments, AAV labeled mid-ventral interneurons and medial motor column motor neurons (**Figure 2F**).

In contrast to the intraspinoventricular injections at NF stage 57, which did not label motor neurons (**Figure S2G-H**), direct injection of AAV into the spinal cord parenchyma at the same stage labeled motor neurons (**Figure S2K-L**), a postmitotic cell type born several stages earlier at NF50-54 ^81^. This difference in the tropism based on delivery method indicates that ventricular administration of AAV is required to label spinal cord temporal cohorts, whereas intraparenchymal administration targets local and postmitotic neurons, as in mice ^55,56,86^.

Across the CNS, in addition to neurons, radial glia were sparsely labeled after AAV and EdU co-injections (**Figure 2B’,G’**). Radial glia were identified by the lack of Elavl3 expression, their elongated morphology, and their hallmark position in the progenitor zone of the spinal cord. This radial glia labeling demonstrates that AAV5 can transduce and drive GFP expression in neural progenitors of both the *Xenopus* spinal cord and brain, suggesting a potential mechanism underlying isochronic cohort labeling (see **Supplementary Discussion**).

These findings indicate that AAV5 transduces neurogenic regions of the tadpole brain and spinal cord, and are consistent with AAV5 functioning as an isochronic cohort marker in the *Xenopus* central nervous system.

### AAVs label isochronic cohorts of developing salamander neurons

The ability of AAVs to label cohorts of neurons generated at the same time during development could result from the transduction of neural progenitors at the ventricle, of neurons that were already born at the time of injection, or both. Distinguishing between these possibilities is necessary to determine whether AAVs can be used as a viral equivalent of EdU for neuronal birthdating experiments.

To address this question, we turned to the dorsal telencephalon or pallium. As previously reported in *Xenopus*, in this brain region, neurons born at different time points are arranged at different distances from the ventricle, with early-born neurons more superficial (further from the ventricle), and late-born neurons deeper (closer to the ventricle) ^78^. This layered distribution is ideal for identifying cohorts of neurons at high temporal resolution. In line with previous *Xenopus* data ^78^, AAV injection in the telencephalic ventricle of *Pleurodeles* early active larvae (stage 41) resulted, three weeks later, in the labeling of neurons confined to a superficial layer of cells, distant from the ventricular zone (**Figure 1G**).

The labeling of superficial neurons after AAV injection could be explained by two alternative hypotheses (**Figure 3A**). AAVs could transduce neural progenitors, so-called radial glia, located at the ventricle, with only cells in their last cell cycle (*i.e.* future neurons) retaining enough viral payload for detectable GFP expression three weeks later. Alternatively, AAV injected in the ventricle could directly transduce neurons that were already post-mitotic at the time of injection; these neurons would then be pushed to the surface by neurons generated later in development. Simultaneous injections of AAVs and EdU can be used to distinguish between these alternatives.

**Figure 3.**
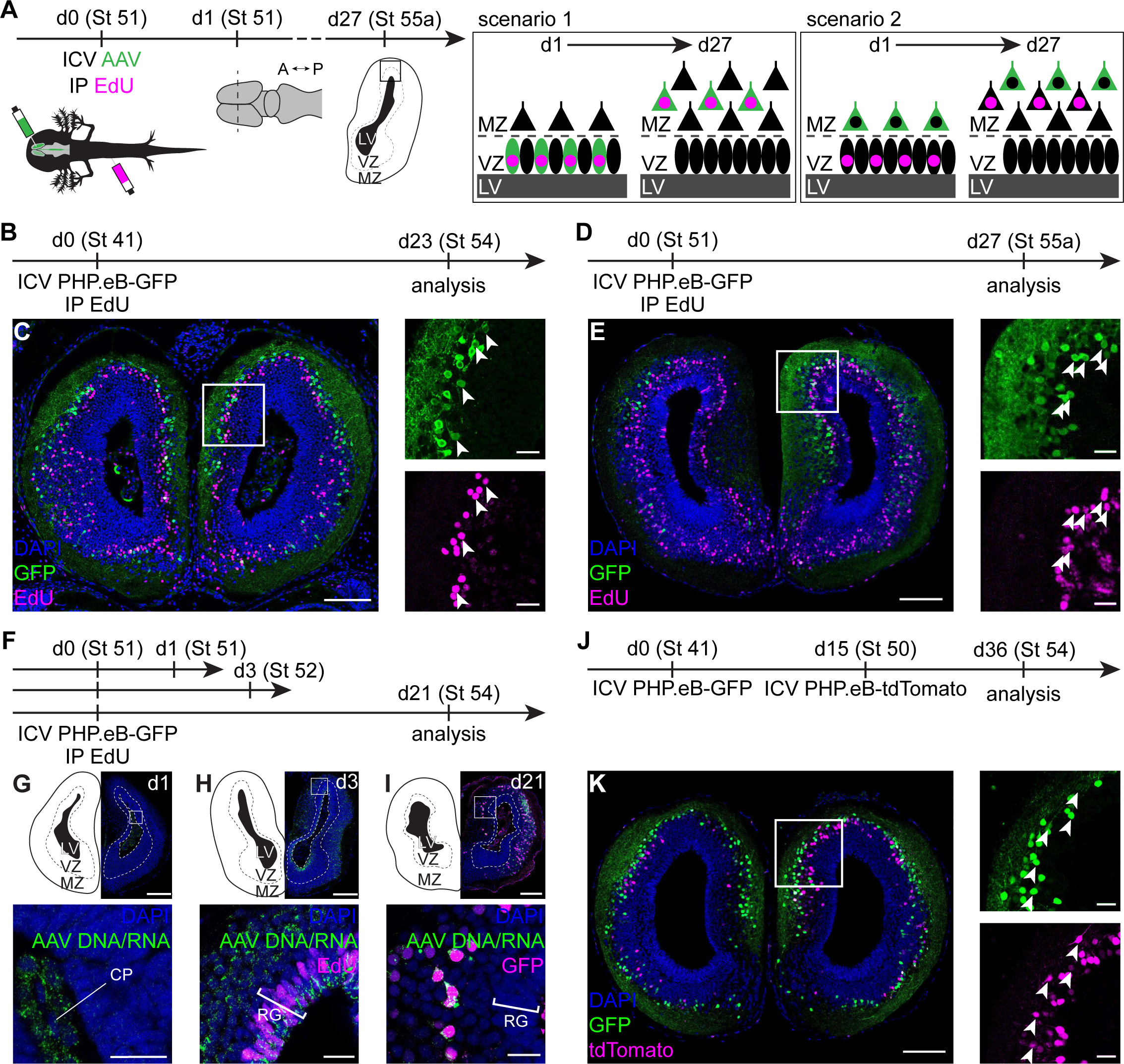
Labeling of different isochronic cohorts of neurons in the *Pleurodeles* telencephalon. **A**. Schematic overview of two possible scenarios of viral labeling. Intracerebroventricular (ICV) injection of AAVs can lead to transduction of proliferating radial glia in the ventricular zone, which would also incorporate EdU if in S phase (scenario 1), or transduction of differentiated neurons in the mantle zone (scenario 2). Three weeks after injection, this would result in EdU+ GFP+ neurons in scenario 1, or differential neuronal labeling in scenario 2. **B**. Schematic overview of the experimental setup: ICV injection of PHP.eB-CAG-GFP and intraperitoneal (IP) injection of EdU in an early-active larva (stage 41), and analysis 23 days later. **C**. Coronal section through the *Pleurodeles* telencephalon showing GFP (green) and EdU (magenta) labeled cells. Box indicates magnified region, arrowheads point towards double-labeled cells. **D**. ICV injection of PHP.eB-CAG-GFP and IP injection of EdU in a late-active larva (St 51), and analysis 27 days later. **E**. Coronal section through the *Pleurodeles* telencephalon showing GFP and EdU labeled cells. Box indicates a magnified region, where white arrowheads point towards double-labeled cells. **F**. ICV injection of PHP.eB-CAG-GFP and IP injection of EdU in a late-active larva, and analysis 1, 3 or 21 days later. **G-I**. Coronal sections through the *Pleurodeles* telencephalon 1, 3 or 21 days after injection, showing presence of viral DNA/RNA (green) in the choroid plexus at day 1 (d1); viral DNA/RNA and EdU (magenta) in radial glia lining the ventricle and neurons in the mantle zone of the telencephalon at d3; and viral DNA/RNA and GFP (magenta) protein in neurons at d21. Boxes indicate magnified regions. **J**. ICV injection of PHP.eB-CAG-GFP in an early-active larva (stage 41) and ICV injection of PHP.eB-CAG-tdTomato in a late-active larva (stage 50); analysis 21 days after the last injection. **K**. Coronal section through the *Pleurodeles* telencephalon showing GFP (green) and tdTomato (magenta) labeled cells. Box indicates magnified region, where arrowheads point towards double-labeled cells. DAPI, blue. Scale bars in overview images and magnifications are 200 μm and 40 μm, respectively. **Abbreviations:** A, anterior; CP; choroid plexus d, day; ICV, intracerebroventricular; IHC, immunohistochemistry; IP, intraperitoneal; ISH, *in situ* hybridization; LV, lateral ventricle; MZ, mantle zone; P, posterior; RG, radial glia; St, stage; VZ, ventricular zone.

To test these hypotheses, we combined intracerebroventricular injection of the AAV-PHP.eB virus with intraperitoneal injection of EdU in stage 41 *Pleurodeles* larvae (**Figure 3B**). Although both labels were incorporated into neurons, the overlap was limited to a small fraction of cells, significantly less than expected if AAVs only transduce radial glia progenitors (see **Supplementary Discussion**). Furthermore, GFP+ and EdU+ cells were found at different distances from the ventricle, supporting the idea that these cells were born at different time points, with AAV-labeled cells born before EdU+ cells. Injections of AAV and EdU in stage 51 larvae yielded the same result, namely the layered distribution of GFP+ and EdU+ cells (**Figure 3B-E**).

To directly test whether AAV-PHP.eB transduces postmitotic neurons, we used *in situ* hybridization chain reaction (HCR) ^87^ with probes that hybridize to parts of the viral genome and also to the viral mRNA. Larvae injected with AAV and EdU were analyzed by HCR at different time points after injection (**Figure 3F**). One day after injection, we detected AAV DNA/RNA in the choroid plexus, indicating rapid clearance of the virus from the ventricle, potentially as a form of immune response ^88^ (**Figure 3G**). Three days after injection, viral DNA/RNA was present in both radial glia in the ventricular zone and in neurons in the mantle zone (**Figure 3H**). The absence of EdU signal in the mantle zone indicates that neurons transduced by AAV were already postmitotic at the time of AAV injection. Three weeks after injection, co-labeling of the viral DNA/RNA and GFP protein was restricted to neurons, whereas viral DNA/RNA and GFP protein were not detected in radial glia cells in the ventricular zone (**Figure 3I**). These results are consistent with the hypothesis that AAV-PHP.eB transduces both radial glia and post-mitotic neurons in *Pleurodeles*, but then results in payload expression only in neurons. The absence of a complete overlap of AAV and EdU labeling also shows that AAV-PHP.eB and EdU cannot be used interchangeably for birthdating studies in the *Pleurodeles* forebrain.

Next, we asked whether AAVs injected in the telencephalic ventricle at a certain developmental time point transduced every neuron already postmitotic at the time of injection, or only a specific subset. Using tandem injections of AAV-PHP.eB expressing either GFP or tdTomato at early (stage 41) and late active (stage 50) larval stages, we found that temporally separated injections of AAV-PHP.eB label largely distinct cohorts of neurons, with a small number of GFP+ tdTomato+ cells (**Figure 3J-K**). Labeled neurons were distributed according to the outside-in sequence of neurogenesis ^78^: GFP neurons (stage 41) were generally more superficial than tdTomato neurons (stage 50). This indicates that intracerebroventricular AAV injections do not label neurons indiscriminately, but target neurons generated in the temporal window around the time of injection. Whether this occurs through transduction of neurons that are already postmitotic but still immature, or some other mechanism, remains to be explored.

The transient detection of AAV DNA/RNA (**Figure 3H**), but not GFP protein (**Figure 3I**), in radial glia indicates that viral genomes are either diluted rapidly during cell proliferation, or that AAV payload transcription or translation are impaired specifically in this cell type ^89^. To distinguish between these two possibilities, we performed intracerebroventricular AAV injections in post-metamorphic juveniles (stage 56) ^65^, in which the majority of radial glia cells have already transitioned to a quiescent state ^12^. Quiescent radial glial cells are called ependymoglial cells (EGCs); these cells re-enter a proliferative state after brain and spinal cord injury, fueling the complete regeneration of these tissues ^7,8,90–92^. Surprisingly, injections of AAV9-CAG-GFP or AAV-PHP.eB-CAG-GFP into the telencephalic ventricle of post-metamorphic animals produced widespread neuronal labeling in the injected hemisphere (**Figure 4A-C** and **Figure S3A-H**), indicating direct transduction of differentiated neurons. To confirm this, we performed HCR for AAV DNA/RNA three days after AAV-PHP.eB and EdU injections, and found AAV DNA/RNA signal both in neurons in the mantle zone and in the EGCs at the ventricle (**Figure 4D-E)**. Labeled neurons were already postmitotic at the time of injection, as indicated by lack of EdU uptake.

**Figure 4.**
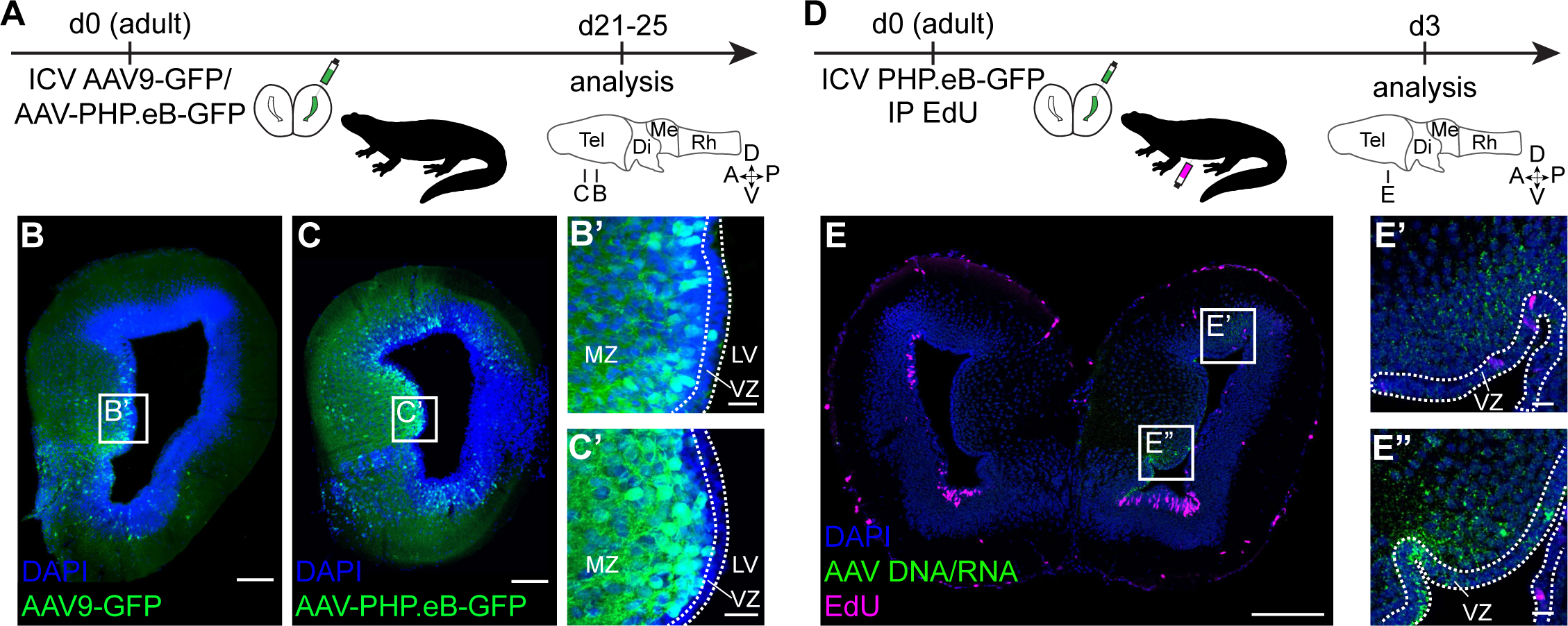
Intracerebroventricular AAV injections produce widespread labeling of neurons but not ependymoglia in the post-metamorphic *Pleurodeles* telencephalon. **A.** Experimental design: intracerebroventricular (ICV) injection of AAV9-CAG-GFP or PHP.eB-CAG-GFP in post-metamorphic *Pleurodeles*, and analysis 21-25 days later. **B-C**. Coronal sections through the *Pleurodeles* telencephalon showing GFP labeled cells (green) in a 5-month old animal injected with AAV9-CAG-GFP (**B**) and in a 3-month old animal injected with AAV-PHP.eB-CAG-GFP (**C**). Boxes indicate magnified regions (**B’-C’**). ICV injection of AAV9 or AAV-PHP.eB leads to transduction of neurons throughout the mantle zone (MZ), and limited transduction of ependymoglia in the ventricular zone (VZ). **D**. Experimental design: ICV injection of PHP.eB-CAG-GFP and IP injection of EdU in a post-metamorphic adult animal, and analysis 3 days later. **E**. Coronal section through the *Pleurodeles* telencephalon 3 days after injection, showing viral DNA/RNA (green) in both radial glia lining the ventricle and neurons in the mantle zone. Viral transduction is not correlated with cell proliferation in the telencephalon of post-metamorphic salamanders, as indicated by lack of overlap with EdU (magenta). Boxes indicate magnified regions (**E’-E’’**). Scale bars in overview images and magnifications are 200 μm and 40 μm, respectively. DAPI, blue. **Abbreviations:** A, anterior; D, dorsal; Di, diencephalon; ICV, intracerebroventricular; IP, intraperitoneal; LV, lateral ventricle; Me, mesencephalon; MZ, mantle zone; P, posterior; Rh, rhombencephalon; Tel, telencephalon; V, ventral; VZ, ventricular zone.

In post-metamorphic animals, AAV9 and AAV-PHP.eB intracerebroventricular injections resulted in extremely sparse GFP protein expression in EGCs, despite that AAV DNA/RNA could be broadly detected in this cell type three days after injection (**Figure 4**), and in line with our results in larvae (**Figure 3I**). Given that electroporation of constructs expressing fluorescent proteins under the control of the CAG promoter successfully labels EGCs (**Figure S3K-L**), the lack of AAV-driven GFP expression in this cell type suggests the presence of a cell-type specific repression of viral payload expression ^93^.

Our experiments thus indicate that intracerebroventricular AAV injections in *Pleurodeles* result in the transduction of isochronic cohorts of neurons in larvae, and ubiquitous but unilateral transduction of neurons in post-metamorphic juveniles. Although the virus also transduces radial glia cells and EGCs, expression of the viral payload in these cell types is impaired after viral entry.

### AAVs transduce neurons in post-metamorphic amphibians

Informed by our results in frog and salamander larvae, we sought to determine whether serotypes that infect the developing nervous system by ventricular injection also infect the mature adult nervous system of amphibians by intraparenchymal injection. Successful injection directly into the CNS parenchyma would confer the ability to target neurons in post-metamorphic animals with tight spatial control, providing a tool to dissect and manipulate mature and functional neural circuits in behaving animals.

### AAVs focally transduce differentiated neurons in adult frogs

To evaluate if AAVs label neurons in a focal region after direct injection into the CNS of adult *Xenopus* frogs, we injected the two most efficient AAV serotypes from tadpole stages – AAV5- and AAVrg-CAG-GFP – into the medial/dorsal pallium of mature adult frogs (1.5 to 2 years old, 2 to 5 cm, 1 to 17 grams), corresponding to post-metamorphic stages 0-5 ^94^. Transduction efficiency was assessed by sectioning and staining brain tissue three weeks after injection, using immunohistochemistry to further amplify GFP expression. Both AAV5 and AAVrg efficiently transduced neurons around the injection site, as visible in coronal and horizontal sections (**Figure 5A-C)**. Labeling efficiency in *Xenopus* was comparable in both small (**Figure 5A-B**) and larger sexually mature (**Figure 5C**) ^95^ frogs, as well as in animals at varying stages of intermediate maturity (**Table S1 and S2**). In contrast, injection of AAV2, AAVrg, AAV1, AAV5, AAV9, or AAV-PHP.eB in the telencephalon (intracerebroventricular injections) of wild-caught adult *Pelophylax bedriagae* did not produce any GFP expression (**Table S1**).

**Figure 5.**
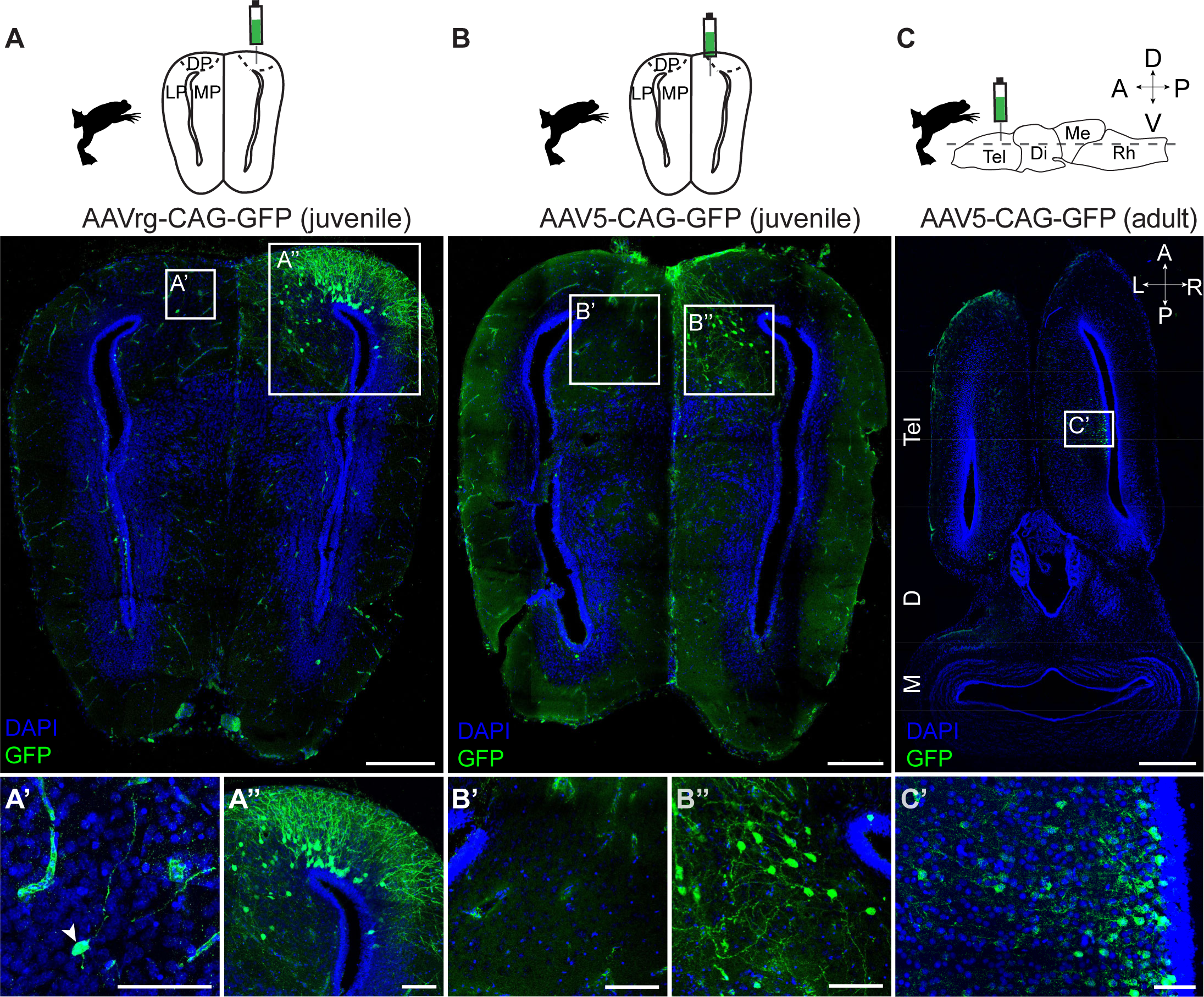
AAVrg and AAV5 transduce neurons in *Xenopus* juveniles and adults. Direct intrapallial injections were performed in post-metamorphic frogs and analyzed three weeks after injection. **A.** AAVrg-CAG-GFP injected into the medial-dorsal pallium transduces sparse neurons in the contralateral pallium (**A’**) and drives strong labeling in the ipsilateral pallium (**A’’**). **B.** AAV5-CAG-GFP injected into the medial pallium (MP) produces local ipsilateral labeling in juvenile frogs. **C.** AAV5-CAG-GFP injected into the MP drives local labeling in larger, sexually mature adult frogs. Sections shown are 40 µm thick and cut either at a coronal (**A-B**) or horizontal (**C**) plane. Scale bars in overview images and magnifications represent 400 µm and 50 µm, respectively. AAV-driven GFP, green. DAPI, blue. **Abbreviations**: A, anterior; D, dorsal; Di, diencephalon; DP, dorsal pallium; LP, lateral pallium; Me, mesencephalon; MP, medial pallium; P, posterior; Rh, rhombencephalon; Tel, telencephalon; V, ventral.

### AAVs focally transduce differentiated neurons in adult salamanders

Next, we asked whether AAV serotypes that transduce the *Pleurodeles* larval brain also transduce older post-metamorphic (5 months to 2 years old, 5 to 20 grams) salamanders, referred to here as adults, after direct injection into the brain parenchyma. We injected AAVs into dorsal pallium, taking care to not inject any virus in the ventricle. Injection of AAV9, AAV-PHP.eB, and AAVrg resulted in GFP reporter expression around the injection site (**Figure 6A-D**). The position and morphology of labeled cells indicated that these viral vectors transduced neurons, characterized by numerous apical dendrites extending toward the pial surface ^96^.

**Figure 6.**
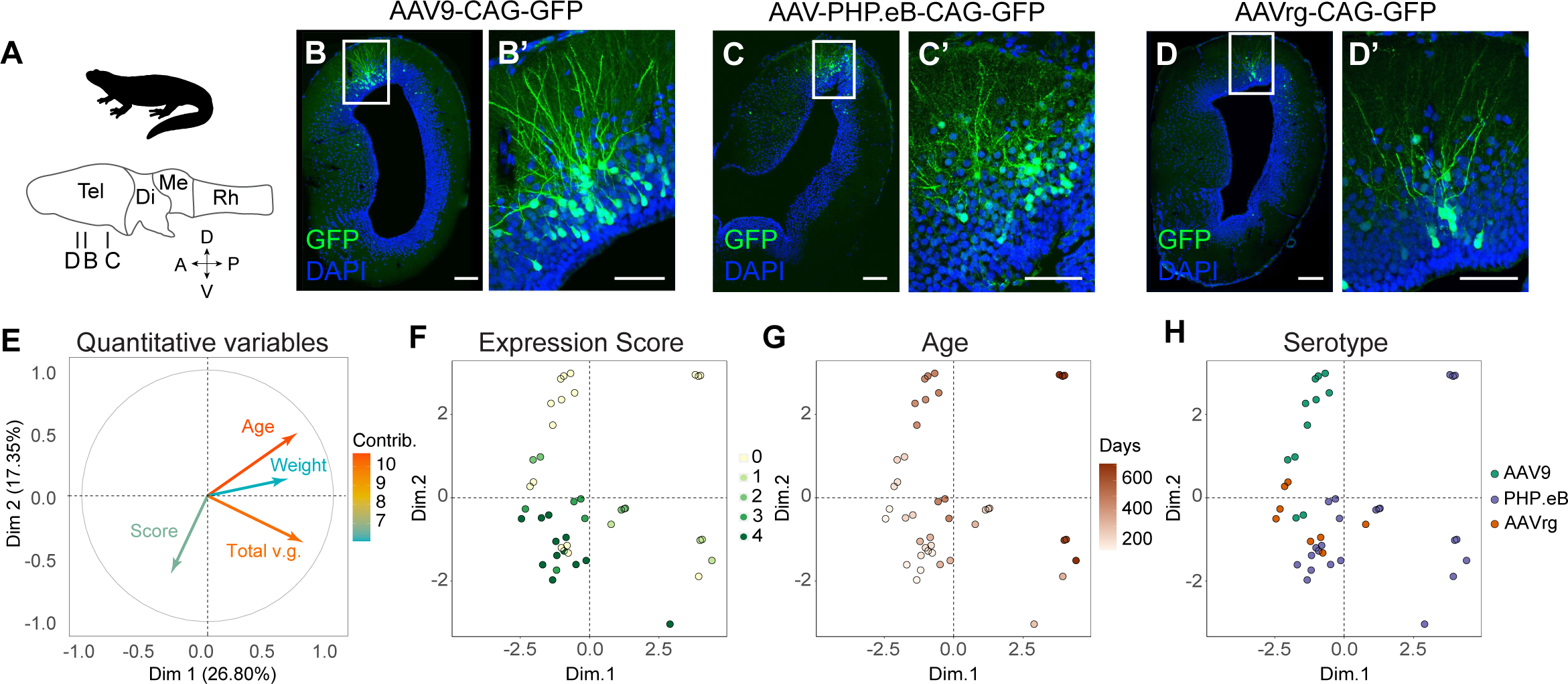
AAV serotype screening by intraparenchymal injections into post-metamorphic *Pleurodeles* adults. **A.** Direct intrapallial injections performed separately in adult (>5 months old) salamanders, with section planes indicated for panels **B-D.** Representative 70 μm thick coronal sections at injection sites for AAV-CAG-GFP (**B**), AAV-PHP.eB-GFP (**C**), and AAVrg-CAG-GFP (**D**). Panels to the left are one entire hemisphere, while panels to the right show detail at the injection site. Scale bars in overview images and magnifications are 200 μm and 100 μm, respectively. AAV-driven GFP, green. DAPI, blue. **E-H.** Injection outcome variability was quantified using Factor Analysis for Mixed Data (FAMD) analysis of 41 individual injections. Both quantitative and categorical variables were analyzed for their contributions to overall variability. Quantitative variables were: animal age and weight, total viral genomes at injection site, and expression score on a scale from 0-4 (see **STAR Methods** for scoring criteria). Categorical variables were: serotype, promoter, reporter, number of constructs injected at one site, manufacturer, and injection site. **E**. Plot showing the contributions of quantitative variables along the first and second FAMD dimensions (x and y axes, respectively). Colors indicate the relative contribution of each quantitative variable to the total variance in the dataset. The x and y coordinates of the arrow tips are correlation coefficients of each variable with the first and second FAMD dimension. **F-H**. Plots showing each individual injection in FAMD space. The two top FAMD dimensions are on the x and y axes, respectively. Each data point is color-coded according to expression score (**F**), animal age (**G**), or serotype injected (**H**). See **Figure S4** and **Table S3** for the full dataset included in FAMD, and additional plotted variables. **Abbreviations:** A, anterior; D, dorsal; Di, diencephalon; Me, mesencephalon; P, posterior; Rh, rhombencephalon; Tel, telencephalon; V, ventral.

In *Pleurodeles*, the results of intraparenchymal injection in post-metamorphic animals were markedly different from the outcomes of intracerebroventricular injections (**Figure 4B,C**). Injections into the telencephalic ventricle produced extensive labeling of neurons throughout the entire telencephalon, which is approx. 3 mm long along the rostro-caudal axis (**Figure S3A-H**). In contrast, after intraparenchymal injections, GFP expression was spatially restricted to neurons within a maximum 200 μm radius around the injection site (data not shown). These results indicate that different injection strategies achieve either widespread or spatially-restricted AAV-driven gene expression in adult salamanders.

### Sources of AAV expression variability in adult salamanders

Unlike AAV injections in larval *Pleurodeles*, in which transduction efficiency was consistent across individuals, AAV injections in adult animals yielded variable results, ranging from hundreds of labeled neurons in some animals to no labeling in others (**Table S1**). We sought to identify the potential sources of such variability.

One potential source is the strength of the promoter driving GFP expression. We tested both CAG and hSyn promoters side by side, to assess the ability of each promoter to transduce salamander cells. Direct comparisons of AAV9-CAG-GFP and AAV9-hSyn-GFP expression after intraparenchymal injections in the pallium showed that CAG drove higher expression levels in neurons in adults (**Figure S4A-C**). However, even though the hSyn promoter was weaker, we could still detect GFP-expressing neurons around the injection site after immunostaining. A comparison of GFP detection with and without immunostaining confirmed that GFP signal amplification is critical to detect transduced neurons when the endogenous fluorescence is weak (**Figure S4D-E)**. These results indicate that promoter choice altered the level of expression but not our ability to detect transduced neurons, and thus could not fully explain the variation we observed.

To query other sources of variation, we scored additional intraparenchymal injections performed in mature adult *Pleurodeles* salamanders, including those in animals of different ages and sizes, with different volumes of virus that varied in their promoter, reporter, and serotype, and with single or dual injections into different brain regions. The full metadata included information on animal weight, single or dual injection, injection site, serotype, promoter, reporter, total viral genomes delivered, and manufacturer, all of which are reported from 41 virus injections in 27 animals (**Table S3**). After immunohistochemistry and confocal imaging, we assigned a cell transduction score ranging from 0 (no expression) to 4 (bright and widespread expression of the reporter, see **STAR Methods**).

To assess the differential contribution of variables in our dataset, we performed a Factor Analysis for Mixed Data (FAMD), a principal component method to explore data with both quantitative and qualitative variables (**Figure 6E-H**) ^97^. As expected, variables including serotype, promoter, and total viral genomes contributed significantly above chance to the variability in the dataset (**Figure S4G-H**). Interestingly, FAMD also indicated that animal age significantly contributed to variability above chance, and that animal age and cell transduction score are broadly anticorrelated, even when all other experimental variables are taken into consideration (**Figure 6F-G, Figure S4I-R**). Linear regression of age against expression score within this dataset confirmed a negative correlation between animal age and transduction efficiency (p<0.005, **Figure S4S**). These results suggest that AAV transduction efficiency in the brain of adult salamanders declines with animal age.

### Retrograde axonal tracing with AAVrg in frogs and salamanders

Of all of the serotypes we screened, only AAVrg efficiently transduced neurons in both frogs and salamanders. AAVrg was generated from an AAV2 capsid library through an *in vivo* directed evolution approach in mice, and selected for its ability to spread retrogradely along axons ^98^. We thus sought to determine whether AAVrg functions as a retrograde tracer in frogs and salamanders.

To evaluate whether AAVrg is retrogradely transported in frogs, we compared the results of direct AAVrg injections with previous knowledge of frog pallial connectivity. The existing literature indicates extensive ipsilateral and sparse contralateral projections in the frog pallium ^99,100^; therefore, after AAV5 or AAVrg intraparenchymal injection, we compared the spread of GFP expression in the ipsilateral and contralateral pallium (**Figure 7A-B and Figure S5**). Direct injection of AAV5-CAG-GFP, a serotype not expected to travel retrogradely, in the medial pallium labeled exclusively neurons in the injected hemisphere (**Figure 5B**), with ∼1mm local spread along the antero-posterior axis (**Figure 7A-B**). In contrast, AAVrg-CAG-GFP injection at the medial-dorsal pallium border resulted in a much wider spread (∼2mm) of viral labeling, suggesting retrograde transport within the pallium. Specifically, AAVrg injections labeled neuronal cell bodies in both the ipsilateral medial, dorsal, and lateral pallium. Moreover, we observed labeled cell bodies in the contralateral medial and dorsal pallium (**Figure 7C-E** and **Figure S5**). This is consistent with previous data in other frog species showing the existence of medial pallium neurons that project to the contralateral pallium ^99,100^. In line with this, we observed GFP-labeled axons passing through the hippocampal commissure (**Figure 7D’’**) and exiting into the contralateral medial pallium near a cluster of labeled cell bodies (**Figure 7D’’’**). In contrast, AAV5-CAG-GFP injected into a similar level of the medial pallium (**Figure 5B**) did not label the commissure or any cells outside the medial pallium injection site. These data support retrograde transport of AAVrg in the *Xenopus* CNS.

**Figure 7.**
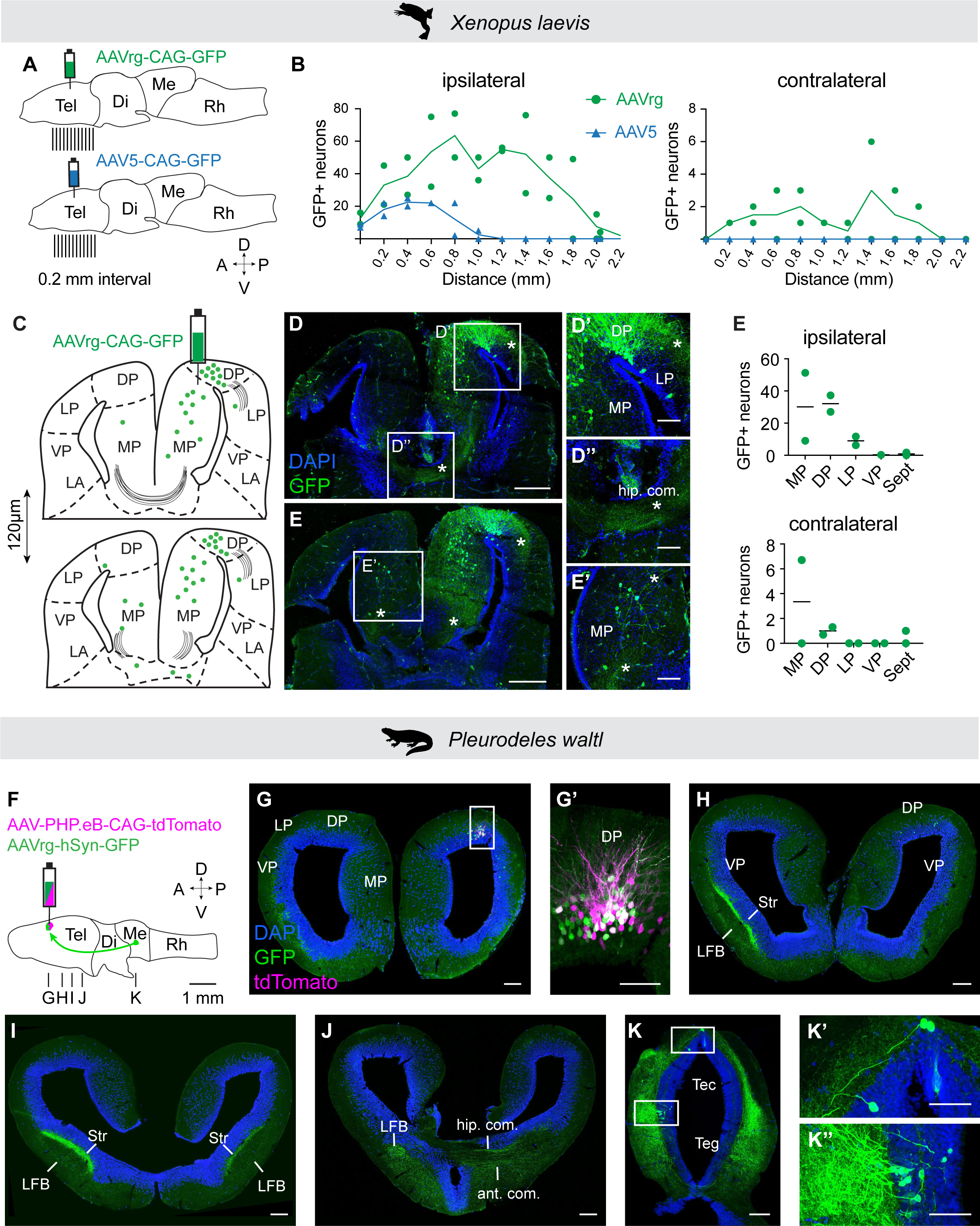
Retrograde tracing of neural circuits with AAVrg in postmetamorphic *Xenopus* and *Pleurodeles.* **A.** Schematic showing injection sites and sectioning planes used for analysis of AAVrg-CAG-GFP and AAV5-CAG-GFP spread after injection into dorsal pallium (DP) of adult *Xenopus*. **B.** Number of transduced neurons across the anteroposterior axis. The numbers of labeled soma on the ipsilateral (left plot) and contralateral side (right plot) are shown for AAVrg (green, n=2) and AAV5 (blue, n=2). Sections amplified for the GFP signal were sampled every 0.2 mm and the labeled soma were counted. The distance shown is from the most anterior to the most posterior labeled cell. **C.** Schematic of two coronal sections from the same brain, with labeling (green dots) and commissures (gray lines) after injection of AAVrg-CAG-GFP into DP/MP, shown in **D.** Coronal sections of pallium showing the local transduction of neurons at the DP injection site and axons to the LP marked by asterisk (**D’**), axonal tracts of the hippocampal commissure marked by asterisk (**D’’**) and traced cell bodies and axons (marked by asterisk) in the contralateral MP (**D’’’**). **E.** Quantification of GFP-positive cells in distinct telencephalic areas after injection of AAVrg-CAG-GFP (n=2). For *Xenopus*, scale bars in overview image and magnifications represent 400 and 50 µm, respectively; AAV-driven GFP, green. DAPI, blue. **F.** Schematic showing injection site and section planes for a dual injection of AAV-PHP.eB-CAG-tdTomato and AAVrg-hSyn-GFP, which were co-injected into the adult *Pleurodeles* dorsal pallium (DP). Arrow indicates axonal projections to DP from the midbrain. **G.** Coronal section showing labeling at the injection site of both tdTomato (magenta) and GFP (green). **G’** Maximum intensity projection of a 25 um stack of 5 images taken at 5 um intervals at the injection site **H-J.** Representative coronal sections along the anterior-posterior axis, showing GFP expression far from the injection site, including strong axonal expression throughout the lateral forebrain bundle (LFB) and commissures. **K.** Retrogradely labeled soma in the midbrain, in both dorsal and ventral tectum (Tec), approx. 3 mm away from the injection site. For *Pleurodeles*, scale bars in overview images and magnifications represent 200 and 100 µm. DAPI, blue. **Abbreviations:** A, anterior; ant. com., anterior commissure; APOA, anterior preoptic areaCeA, central amygdala; D, dorsal; Di, diencephalon; DP, dorsal pallium; hip. com, hippocampal commissure; LA, lateral amygdala; LFB, lateral forebrain bundle; LP, lateral pallium; Me, mesencephalon; MP, medial pallium; P, posterior; Rh, rhombencephalon; Sept, septum; Str, striatal neuropil; Tec, tectum; Tel, telencephalon; Teg, tegmentum; V, ventral; VP, ventral pallium.

To determine whether AAVrg can be used for retrograde tracing in salamanders, we co-injected AAVrg-hSyn-GFP and AAV-PHP.eB-CAG-tdTomato in the most lateral portion of the dorsal pallium of adult *Pleurodeles*. AAV-PHP.eB was not expected to travel retrogradely, and served as an injection control. We observed tdTomato expressing cells in the pallium exclusively at the injection site three weeks post injection (**Figure 7F-G**). AAVrg-driven GFP fluorescence, in contrast, was much more widespread. We observed both GFP+ neuronal cell bodies around the injection site, and GFP+ axonal tracts in the ipsilateral and contralateral striatal neuropil and lateral forebrain bundle (**Figure 7H-I**). Decussating GFP+ fibers were visible in the anterior and hippocampal commissures (**Figure 7J**). In more posterior brain sections, GFP+ axons could be followed throughout the diencephalon and midbrain. In the midbrain, GFP-expressing neuronal cell bodies were observed in multiple areas, including the ventral mesencephalic tectum, about 3 mm away from the injection site (**Figure 7K**). The pattern of GFP expression indicates that AAVrg was taken up by axonal terminals at the injection site and traveled retrogradely to reach the somata of dorsal pallium-projecting neurons in the midbrain. These findings not only show that AAVrg is a retrograde tracer in salamanders, but also corroborate the existence of direct projections from the midbrain to the dorsal pallium, as initially indicated by experiments with dextran tracers in *Pleurodeles* ^16^.

Our results thus demonstrate that AAVrg is a powerful viral tool for neuronal labeling and axonal tracing in salamanders and frogs.

## Discussion

AAV vectors are a workhorse of modern neuroscience research in rodents ^39,101^ and hold great promise for gene therapy in humans ^102^. Despite having significant potential for research on lesser-studied species not amenable to transgenesis, broader use of AAVs has been limited. These viral vectors have only recently been employed in neuroscience applications in non-human primates ^103,104^, birds ^105–108^, and reptiles ^109^. Previous experiments in zebrafish ^110^ and frogs ^42^ either did not find, or found negligible, AAV transduction in the brain. Here, we demonstrate that AAVs do infect the CNS of three distantly-related species of amphibians. Our findings provide a roadmap for AAV screening in new species, reveal new aspects of AAV biology, and open exciting opportunities for neuroscience research using amphibian species.

### Screening the AAV toolbox in new vertebrate species

Four key elements of our experimental design proved essential for the success of our AAV screen. First, we started from a broad set of serotypes, without assuming that a serotype that works in one amphibian species would also work in others. Indeed, of all serotypes tested, we found none were pan-amphibian in their neuron transduction: AAV5 transduces both *Pelophylax* and *Xenopus* frogs, but its transduction efficiency is extremely low in *Pleurodeles* salamanders; AAV-PHP.eB transduces salamanders and only *Pelophylax* but not *Xenopus* frogs; and AAVrg transduces salamanders and *Xenopus* but not *Pelophylax* frogs. AAV tropism can thus vary drastically even in species belonging to the same order. When transduction was successful, the extent of labeling by each serotype also varied, with the most limited expression observed in *Pelophylax* and perhaps resulting from the high genetic variability, or variation in immune status, of wild-caught animals. We conclude that an unbiased set of serotypes is the ideal starting point for screening AAVs in new species.

Second, the delivery strategy matters. We observed distinct populations of neurons labeled by intraventricular versus intraparenchymal injection. Intraventricular injections in larvae labeled developing neurons along the entire rostro-caudal axis of the brain or spinal cord, targeting populations that are born around the time of injection, as in mammals ^56^. In contrast, intraparenchymal injections labeled only neurons around the injection site, with the exception of AAVrg which captured both the cell bodies within and neurons projecting to the injection site. Notably, the speed and volume of our intraparenchymal injections in adults differed markedly from previous experiments in frogs ^42^, indicating slow and low-volume virus delivery is essential for transduction.

Third, our ability to detect transduced neurons was dramatically enhanced by a strong and ubiquitous promoter, and by immunohistochemical signal amplification. Although we found that both the CAG and hSyn promoters drive reporter expression in neurons, CAG-driven expression was brighter and easier to detect. Nevertheless, signal amplification by immunohistochemistry ensured that, with successful serotypes, some labeling could always be observed with either promoter or condition tested.

Finally, when designing a screen, life stage is also an important factor to consider. Here, our observations are reassuring: we found that AAVs that efficiently transduce tissue during larval and tadpole development will also work in adults. In *Xenopus*, labeling efficiency at four stages is comparable. However, in older adult *Pleurodeles*, transduction efficiency seems to decrease with age. This observation motivates a two-tiered screening design: first targeting a large number of neurons and brain regions in developing animals, which are often more accessible, and then, verifying that the top candidates from the developmental screen transduce adult animals.

### Lessons on AAV biology from Amphibians

Our experiments in amphibians also suggest new aspects of AAV biology, which inform the design and optimization of AAV vectors as tools for gene delivery in new organisms.

The lack of demonstrated success of AAVs in aquatic vertebrates ^110^ until now seemed to suggest that AAV vectors are functional only in terrestrial animals or only in endotherms. Indeed, all commercially-available vectors were derived from viruses that use mammalian species as natural hosts ^49^. However, our experiments show that AAV transduction is efficient also in aquatic ectotherms living at lower temperatures: *Xenopus* at 18-22°C, *Pelophylax* at approx. 20-24°C in the wild during breeding season, and *Pleurodeles* at 20-23°C. These results extend further the permissive temperature range from recent AAV data in turtles and lizards, which live at ∼ 24°C, but can reach over 40°C during basking ^109^. Our observations imply that body temperature is not a limiting factor for the application of mammalian-derived AAV vectors to a larger set of vertebrate species. This raises the potential that further testing in other aquatic species, such as fish ^110^, may be warranted.

Our comparisons between species, life stages, and cell types revealed differences between AAV serotypes that may underlie the biological mechanisms governing uptake and expression of these viral vectors. Although the rules for AAV cell attachment and entry are not fully understood, it is clear that each serotype uses a specific combination of cell surface glycan and protein receptors ^111^. Our results suggest that these complex combinations of glycans and receptors are highly evolvable across species. Both AAV-PHP.eB and its parent virus AAV9, which in mammals use galactose, Laminin Receptor (LamR), KIAA0319L (AAVR), and GPR108 for cell entry and intracellular trafficking ^112^, worked in *Pleurodeles* but not in *Xenopus*. Conversely, AAV5, the most distantly related to all other AAV serotypes, was the most efficient virus in *Xenopus* and among the least efficient tested in *Pleurodeles.* In mammals, AAV5 uses sialic acid, platelet-derived growth factor receptor (PDGFR), and AAVR receptors ^113^ while notably being GPR108-independent ^114,115^. While the genes for AAVR, LamR and PDGFR are conserved across amphibians, there is no evidence for the presence of GPR108 receptor gene in frog genomes ^116,117^. Thus, the absence of this receptor may indicate why AAV5, and not AAV9 or AAV-PHP.eB, is the most successful frog serotype. Other differences in the performance of serotypes between species may be explained by highly evolvable residues on these proteins, or their recruitment of entirely different combinations of glycan and protein receptors. Our results extend observations in mammals, where comparisons across mouse strains and species of non-human primates have revealed substantial differences of AAV tropism ^103^.

Interestingly, AAVrg was the only serotype able to transduce both salamander and frog neurons with high efficiency, and exhibited retrograde transport, as in mice and lizards ^98,109^. AAVrg was derived from AAV2, which does not efficiently infect any of the species tested here, and its capsid harbors a 10 amino acid insertion in the heparan sulfate binding loop that weakens its heparin affinity, plus two additional mutations ^98^. It has been suggested that this insertion might be critical for the interaction of the viral capsid with the retrograde transport machinery ^118^. The ability of AAVrg to travel retrogradely in several species of mammals ^98,119^, in reptiles ^109^, and in amphibians - though not in birds ^106^ - suggests that the mechanisms of capsid interaction with the retrograde transport machinery might be evolutionarily conserved.

After AAV cell entry, interactions between viral particles and intracellular machinery are crucial for successful payload expression. Some of these interactions might be cell-type specific, determining AAV cellular tropism. In *Pleurodeles*, AAV-PHP.eB drives expression in neurons, but fails to express protein in radial glia and EGCs, even though viral nucleic acids can be detected in these cell types. This could be due to the translational repression of viral mRNA in these glial types. Alternatively, the AAV genome might be epigenetically silenced in radial glia ^89^, or its translocation to the nucleus might be prevented, followed by degradation of the viral DNA, as is suggested for mouse microglia ^120^. Interestingly, radial glia and EGCs are the glial cell types most closely related to mammalian astrocytes, which are transduced by AAV-PHP.eB with high efficiency ^121^. Taken together, our observations indicate that differences in AAV tropism across species may depend not only on the evolutionary divergence of cell surface receptors, but also on species-specific and even cell-type specific differences of intracellular processing of viral particles.

It is conceivable that the complex molecular pathway from AAV vector internalization to transgene expression can be not only cell-type specific, but also dynamic across the lifetime of an organism. In *Pleurodeles*, we observed a decrease of transduction efficiency specifically correlated with animal age. Age-dependent changes of AAV tropism have also been observed in rodents ^55,122^. These observations suggest that age-dependent gene expression changes of the brain tissue, immune system, or even the sexual maturation may impact AAV performance.

### Capturing isochronic cohorts of neurons by developmental time using AAVs

In invertebrate and vertebrate nervous systems, the position, connectivity, and even functional properties of a neuron are often correlated with its developmental history. Neuronal birth order is linked to the anatomical organization of brain regions ^123,124^, cell identity specification ^125^, and the establishment of circuit connectivity ^56,126,127^. Our study demonstrates that intraventricular injection of AAVs in the brain and spinal cord of developing amphibians results in reliable transduction of neuronal populations that develop around the time of injection, termed isochronic cohorts. AAVs thus provide a tool to access these isochronic units in amphibians and evaluate how relationships between development, anatomy, circuit assembly, and function develop and may have evolved.

The ability to label and target isochronic cohorts will provide new insights on long-standing questions in developmental neurobiology - for example, whether and how neural circuits are remodeled during metamorphosis as amphibians transition from aquatic swimming to walking ^128–130^. AAVs will allow selective visualization and comparison of the projection patterns of neurons generated at different stages around metamorphosis. Furthermore, manipulating neuronal activity in different isochronic cohorts may clarify whether early larval circuits are necessary for the proper development and function of late adult circuits.

Comparing isochronic cohorts of neurons across species will also shed light on the evolution of brain regions and neural circuits. Our results indicate that, like in mammals, complex brain regions such as the pallium and the optic tectum are built by adding new neurons in precise temporal patterns (layers in the pallium or cortex, wedges in the optic tectum or superior colliculus, and dorso-ventral domains in the spinal cord) also in amphibians. AAVs, in combination with scRNAseq or spatial transcriptomics, will enable a comparison of the molecular identities and projections of neurons generated in the same relative order in different species ^131^. Going beyond individual brain regions, labeling and manipulating isochronic cohorts of neurons in the entire CNS may be used to investigate the relationships between neuronal birthdate and connectivity, and to test, for example, the hypothesis that neurons that are born together wire together ^127^. In an evolutionary framework, this may reveal general principles for the assembly of neural circuits in the brain and spinal cord.

### Tracing and manipulation of neural circuits using AAVs

The conserved ability of AAVrg to travel in a retrograde manner in both *Xenopus* and *Pleurodeles* has significant potential for the characterization and manipulation of neural circuits in amphibians.

Neural circuit analysis in amphibians is critical for establishing a complete picture of vertebrate brain evolution and for understanding the adaptations associated with life on land. Yet, several aspects of the amphibian brain remain enigmatic. The lack of conspicuous neuroanatomical landmarks, such as clearly distinguishable cytoarchitectural boundaries, presents challenges to the identification and comparison of brain nuclei. This is further exacerbated by the small size of the brain and regions therein. More than in any other vertebrate clade, molecular data have been essential to clearly delineate brain regions, leading to the discovery of features, such as layers in the pallium, that were thought to be secondarily lost in amphibians ^16,132^.

Similar to how molecular tools revolutionized our understanding of brain regions, the availability of AAVs for axonal tracing experiments will dramatically improve the quality, scale, and resolution of connectivity data for amphibians, as already indicated by our initial experiments. Our AAVrg *Xenopus* experiments confirm the existence of reciprocal connections between the medial pallia, which were only reported in a distantly-related frog species ^99,100^. In *Pleurodeles*, we find a direct projection from the ventral tectum to the dorsal pallium that was largely uncharacterized ^132,133^. These dorsal pallium-projecting neurons might be homologous to parts of the torus semicircularis (ventral part of tectum) in zebrafish, which also sends direct ipsi- and contralateral projections to the pallium through the lateral forebrain bundle ^134^. The input-output connectivity of these neurons in fish and salamander suggest a role in processing mechanosensory inputs from the lateral line ^135^, possibly explaining the absence of this direct midbrain-pallium projection in adult anurans and amniotes ^136^, who have lost the lateral line.

These examples indicate the importance of high-resolution tracing data for the accurate characterization of the amphibian brain and inferences of brain evolution. Furthermore, AAV-based tracing experiments not only yield data with high signal-to-noise, but are also compatible with whole-brain clearing and gene expression profiling ^137,138^, and can be further tailored to specific neuron types by using constructs with cell-type specific promoters or enhancers ^105,139^. The integration of these tools will produce multimodal, high-resolution atlases for the amphibian brain, complementing those available for fish and mammals ^137,140–143^.

Beyond comparative neuroanatomy, AAVs open the door to the genetic manipulation of neural circuits in amphibians. One application of particular interest would be to express optogenetic actuators in specific populations of cells, as defined by their projection identities. Given the transparency of *Xenopus* and *Pleurodeles* skin at earlier stages and the existence of non-pigmented lines, the use of light to non-invasively stimulate or inhibit subpopulations of neurons and probe their role in behavior is now feasible in these species. Additionally, given that we were able to identify other serotypes that do not have this retrograde transport activity, it will be possible to employ even more fine-tuned targeting of CNS populations, using Cre-based targeting strategies commonly used in mammalian systems to target subsets of projection neurons ^98^.

Our study establishes an initial AAV toolkit that may be further optimized and extended by engineering ^98,144,145^ to other amphibian species, thus empowering the dissection of neural circuit assembly, structure, and function in this important taxonomic group. This toolkit opens new avenues for the functional study of the amphibian CNS, increasing our understanding of the diversity of developmental processes and neural circuits that have arisen during vertebrate evolution.

## Acknowledgements

We would like to extend our thanks to members of the Sweeney, Tosches, Shein-Idelson, Yamaguchi, Kelley, and Cline Labs for their contributions to this project, discussion and support. We additionally thank the Beckman Institute Clover Center and Viviana Gradinaru (Caltech), Kimberly Ritola (UNC NeuroTools), Flavia Gama Gomez Leite (ISTA Viral Core), and Hüseyin Cihan Önal (Shigemoto Group, ISTA) for their consultation and assistance regarding AAVs, as well as Andras Simon and Alberto Joven for feedback and discussions on AAVs in *Pleurodeles*. To do these experiments, we have also benefited from the tremendous support of our animal care and imaging facilities at our respective institutions, as well as the amphibian stock centers (National Xenopus Resource Center, European Xenopus Resource Center, Xenopus Express) and our funding sources: U.S. National Science Foundation (NSF) Grant Number IOS 2110086 (D.B.K., L.B.S., M.A.T., A.Y., and H.T.C.); United States-Israel Binational Science Foundation (BSF) Grant Number 2020702 (M.S.-I.); NSF Award Number 1645105 (G.J.G., M.E.H.); FTI Strategy Lower Austria Dissertation Grant Number FTI21-D-046 (D.V.); Horizon Europe ERC Starting Grant Number 101041551 (L.B.S.); NIH grant number R35GM146973 (M.A.T.); Rita Allen Foundation award number GA_032522_FE (M.A.T.); European Molecular Biology Organization Long-Term Fellowship ALTF 874-2021 (A.D.); National Science Foundation Graduate Research Fellowship DGE 2036197 (E.C.J.B.); NIH grant number P40OD010997 (M.E.H).

## Author Contributions

M.A.T. and L.B.S. led and coordinated the project. E.C.B.J., D.V., A.D., H.T.C., T.F.S., D.B.K., A.Y., M.S-I., M.A.T., and L.B.S devised the project and implemented experiments. E.C.B.J., A.D., J.W., A.O.G., and M.A.T. collected the *Pleurodeles* data; D.V., A.L.N., G.I., R.C.A., F.B., F.A.T., A.Y and L.B.S. collected the *Xenopus* data; N.Z., A.S., and M.S-I. collected the *Pelophylax* data. G.J.G. and M.E.H. provided the *Xenopus* cell line. E.C.B.J., D.V., A.D. N.Z., G.I., M.A.T., and L.B.S analyzed the data. E.C.B.J., D.V., A.D., M.A.T., and L.B.S. wrote the manuscript. H.T.C., T.F.S., D.B.K., A.Y., and M.S-I. edited the manuscript and provided critical feedback on the project.

## STAR Methods

## RESOURCE AVAILABILITY

### Lead contact

Further information and requests for resources and reagents should be directed to and will be fulfilled by the lead contact, Lora B. Sweeney.

### Materials availability

This study did not generate new unique reagents.

### Data and code availability

Any additional information required to reanalyze the data reported in this paper is available from the lead contact upon request.

## KEY RESOURCES TABLE

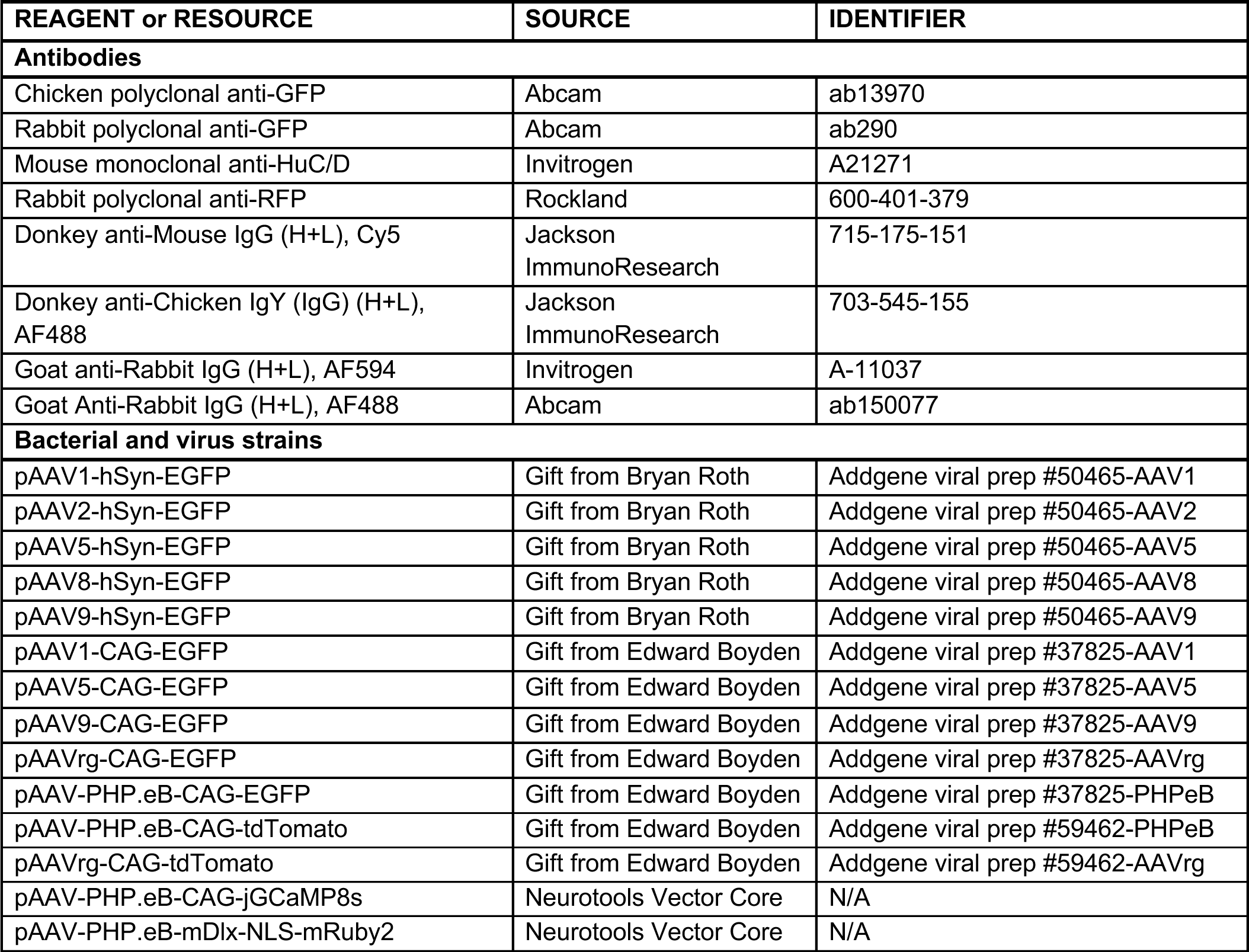
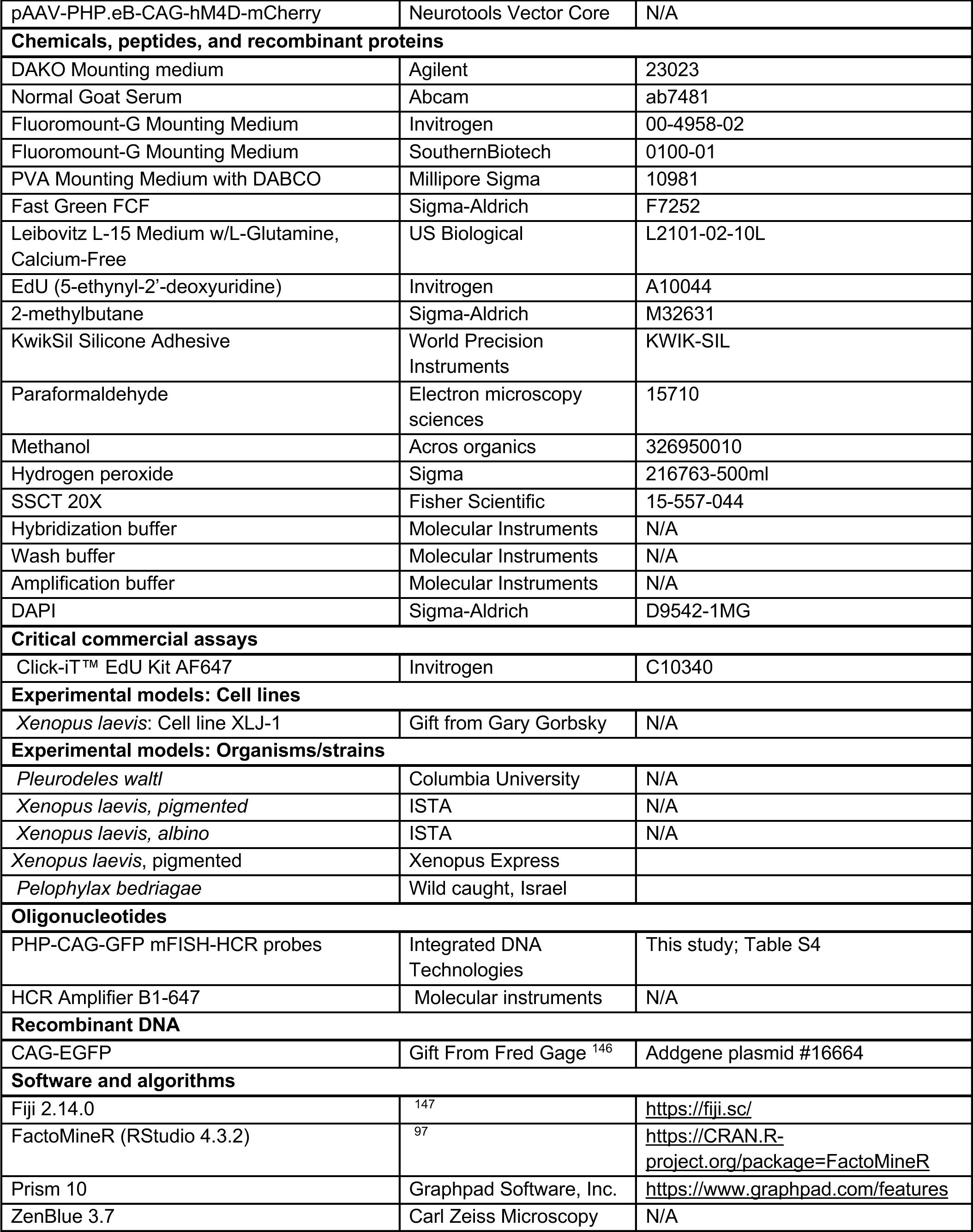
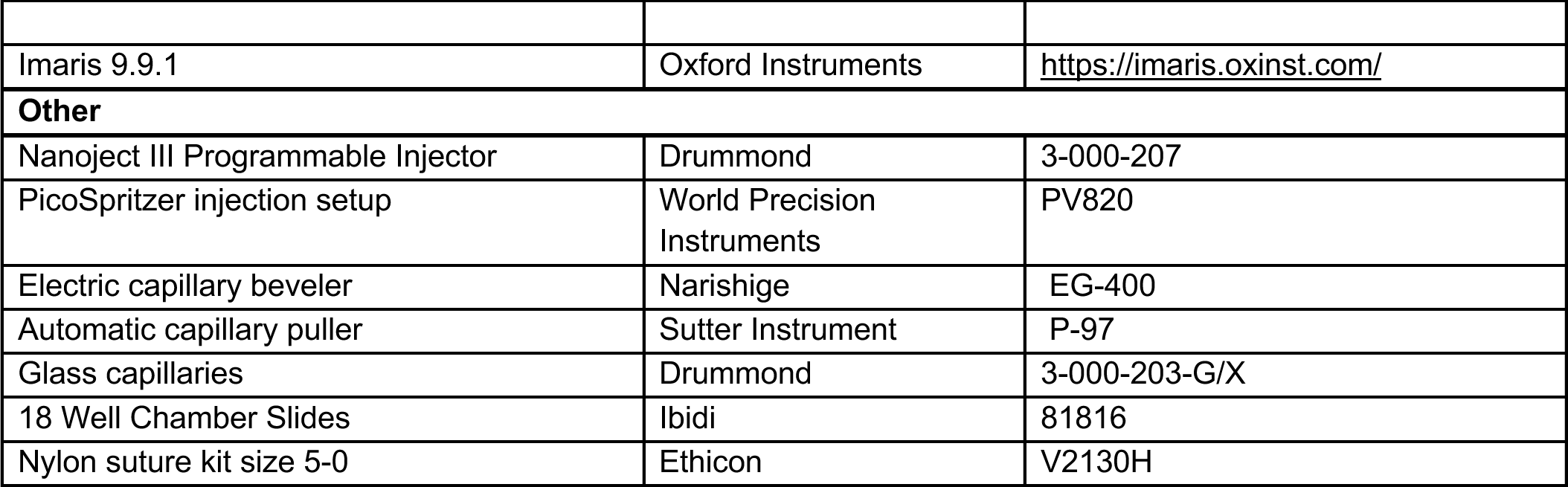

## EXPERIMENTAL MODEL AND STUDY PARTICIPANT DETAILS

*Pleurodeles waltl* were obtained from breeding colonies established at Columbia University. Animals were maintained in an aquatics facility at 20-25 °C under a 12L:12D cycle ^22^. All experiments were conducted in accordance with the NIH guidelines and with the approval of the Columbia University Institutional Animal Care and Use Committee (IACUC protocol AC-AABF2564). Experiments were performed with adult (4.5-24 months) male and female salamanders, and stage 41 and 50/51 larvae (staged according to ^65^).

Albino and pigmented *Xenopus laevis* tadpoles and frogs were either bred and raised at the Institute of Science and Technology Austria (ISTA), or obtained from Xenopus Express and raised at the University of Utah. They were maintained in frog facilities with the water temperature of 18-22 °C under a 12L:12D cycle. For larval serotype screening, albino and pigmented tadpoles NF stage 40-60 (staged according to ^68^) were used for experiments. For juvenile serotype screening, 14 pigmented postmetamorphic animals, stages 0 to 5 (determined according to ^94^), were used. Their mass ranged from 1.08 to 17.48g, and snout-vent length ranged from 2.2 to 5.3cm (for individual animal measurements see **Table S2**). All experiments were performed in accordance with the local ethics committee guidelines and the University of Utah Institutional Animal Care and Use Committee (protocol number 2020-0.762.370, 2022-0.137.228, 2024-0.019.606, and

00001947).

*Pelophylax bedriagae* adult frogs and tadpoles were caught in the wild and maintained in a frog facility at 24 °C under a 12L:12D cycle. Tadpoles stage 25-27 (41) were used for experiments. All animal procedures were performed in accordance with the guidelines of the Tel Aviv University ethical committee (animal license TAU-LS-IL-2112-19-4). Animals were collected under the Israel Nature And National Parks Protection Authority approval (2021/42935).

## METHOD DETAILS

### Xenopus cell culture AAV screening

The XLJ-1 cell line was generated from neurula stage *X. laevis* embryos using methods similar to those previously used to create *Xenopus tropicalis* cell lines ^148^. Cells were grown in 70% calcium-free L15 medium (L2101-02, US Biologicals) with 10% fetal bovine serum and 1% penicillin/streptomycin. They were sparsely seeded into 18 Well Chamber Slides (81816, Ibidi) coated with ibiTreat to promote cell adherence. Twenty-four hours later, after the cells attached to the bottom of the dish, the medium was switched to antibiotic-free medium for 2 hours. 5 μL of viral prep was then added to the first well containing 200 μL. 100 μL of dilution was then carried over to the adjacent well in a repeated manner to create a serial dilution (0.0008, 0.00015, 0.0031, 0.00625, 0.0125, and 0.025 μL virus/μL media) within the entire row. The medium was changed after 12 hours, and GFP expression was monitored and imaged every 24 hours using a cell culture microscope (Leica). At the end of the experiment, the cells were fixed with 4% paraformaldehyde for 20 minutes, washed thrice with PBS and imaged on an LSM800 confocal microscope (Zeiss).

### Intracerebroventricular, intraspinoventricular, and intraparenchymal injection in Xenopus tadpoles

*Xenopus laevis* tadpoles stage NF40-60 were anesthetized in 0.01% MS-222 and placed on a piece of gauze in a plastic Petri dish. FastGreen was mixed with AAV prep received from Addgene at a ratio of 1:5 for visualization of injection. Glass capillaries (3-000-203-G/X, Drummond) were pulled at a capillary puller (P-97, Sutter Instrument), their tips trimmed using a fine forceps and beveled at 30 degrees using an electric capillary beveler (EG-400, Narishige). The virus mix was loaded into glass capillaries right before injection using a GelLoader Pipette tip. The capillary was mounted onto a PicoSpritzer injection setup (PV820, WPI) using a piece of tubing while ensuring proper air seal. The PicoSpritzer was switched to gated injection mode. The drop size was normalized using the axioscope eyepiece scale at 2x zoom, setting it to 4-5 lines, approximately 10 nL per drop (per manufacturer’s conversion chart). Using a micromanipulator, the needle was inserted into the midbrain ventricle through the skin at the level of midline, into the spinal canal through the dorsal trunk muscle, or into the spinal parenchyma depending on the type of injection. The animals were injected with 4 pumps of virus mix equal to approx. 40-60 nL and the successful injection was confirmed by green coloring of the ventricular space. The injected tadpoles were then returned to a tank with fresh 0.1xMMR solution. 21 days after injection, the animals were sacrificed, dissected, and fixed for 1.5 hours on ice in 4% PFA/PB. The fixed tissue was washed 3 times with ice cold 1xPBS and cryoprotected shaking overnight at 4 °C in 15% sucrose-PBS solution complemented with 8% cold fish skin gelatine. The following day, the tissue was lightly dried, embedded in cryomolds filled with tissue freezing medium and frozen in 2-methylbutane (M32631-2.5L, Sigma-Aldrich) on dry ice. 40 μm cryosections were cut at a cryostat (Bright). Sections were imaged on an LSM800 confocal microscope (Zeiss), each had 30-40 optical planes with 1.5 μm steps. The images were stitched using ImarisStitcher, and image processing including contrast adjustment, size filtering (in case of autofluorescent debris), and maximum intensity projection, was done in Fiji.

### Intracerebroventricular injection in Pleurodeles larvae

Larvae were deeply anesthetized in 0.02% MS-222 and stabilized in a Sylgard mold. FastGreen was added to AAV aliquots (as provided by AddGene) at a ratio of 1:6 for visualization of the injection site. Glass capillary needles were pulled and then broken open using fine forceps (needle tip diameter ranging 5-15 um), backfilled with mineral oil, and then connected to a Nanoject III injection system (Drummond) and filled with AAV solution. The needle was inserted through the skin, in the lateral telencephalic ventricle of the right hemisphere and 80 nL of virus solution at stage 41, and/or 120 nL at stage 51 was pressure injected (5 nL/sec). For serotype testing, larvae were sacrificed 23 days later and fixed overnight at 4 °C in 4% PFA/PBS. Larvae were then washed in PBS, cryoprotected in 30% sucrose/PBS and embedded in Tissue-Tek OCT compound (Sakura). 20 μm coronal sections were cut on a cryostat and mounted on glass slides. For detection of the viral DNA/RNA, larvae were perfused 1, 3 or 21 days after injection with PBS-DEPC, brains were dissected and fixed overnight at 4 °C in 4% PFA/PBS-DEPC. Brains were then dehydrated (40%, 60%, 80%, 100%, 100% methanol in PBS-DEPC, 15 min each) and incubated in 100% DCM overnight at RT, washed in 100% methanol and stored at −20°C until further processing.

### Intracerebroventricular injection in Pelophylax bedriagae tadpoles

Tadpoles were anesthetized in 0.05% MS-222 and placed on a piece of gauze in a plastic Petri dish. AAV viral vectors mixed with FastGreen for visualization (ratio 1:5) were injected using a PicoSpritzer (Cat. num. PV820, WPI) through pulled (P-30 puller, Sutter Instrument) glass capillaries (Cat. num. 3-000-203-G/X, Drummond) trimmed at their end. Injections were into the rostral brain ventricle. Injections contained ∼ 40 uL (injected in 4 puffs) and were confirmed by green coloring of the ventricular space. After injection, tadpoles were returned to their tank and sacrificed after 21 days. Brains were dissected and fixed for 4-12 hours at 4°C in 4% PFA/PBS. The fixed tissue was washed 3 times with 1xPBS and cryoprotected overnight at 4 °C in 15% sucrose-PBS solution complemented with 8% cold fish skin gelatine. The following day, the tissue was lightly dried, embedded in cryomolds filled with tissue freezing medium and frozen in 2-methylbutane (Cat. num. M32631-2.5L, Sigma-Aldrich) on dry ice. 40 μm cryosections were cut at a cryostat (Leica CM1950) and stained using immunohistochemistry.

### Intrapallial injections in Xenopus juveniles and adults

Pigmented juvenile *Xenopus* frogs were deeply anesthetized and wrapped in MS-222 soaked gauze to keep them moist. The region of interest was exposed, the craniotomy was performed using a dental drill (Foredom) equipped with a 0.75 mm carbide drill bit (Stoelting), and the meninges were removed using fine forceps. FastGreen was mixed with AAV prep received from Addgene at a ratio of 1:5 for the visualization of the injection site. Glass capillaries were pulled at a capillary puller (P-97, Sutter), their tips trimmed using a fine forceps and beveled at 30 degrees using an electric beveler (EG-400, Narishige). The beveled capillaries were backfilled with mineral oil, mounted to a Nanoject III injection setup (Drummond), and loaded with virus mix. 100 nL of AAV solution was then pressure injected into dorsal telencephalon at an injection rate of 1 nL/12 sec or 1nL/sec, for a total injection time of approx. 20 min. After the injection, the needle was left in the tissue for 5 minutes to prevent leakage. The skin was sutured using nylon suture kit size 5-0 (V2130H, Ethicon). 21 days after injection, the animals were sacrificed, the brains dissected out, and fixed overnight at 4 °C in 4% PFA/PB. The fixed tissue was washed 3 times with ice cold 1xPBS and cryoprotected shaking overnight at 4 °C in 15% sucrose-PBS solution complemented with 8% cold fish skin gelatin. The following day, the tissue was lightly dried, embedded in cryomolds filled with tissue freezing medium and frozen in 2-methylbutane (M32631-2.5L, Sigma-Aldrich) on dry ice. 40 μm cryosections were cut at a cryostat (Bright). Sections were imaged on an LSM800 confocal microscope (Zeiss), each had 30-40 optical planes with 1.5 μm steps. The images were stitched using ImarisStitcher, and image processing including contrast adjustment, size filtering (in case of autofluorescent debris), and maximum intensity projection, was done in Fiji. The expression strength of adult injections was scored in **Table S2**, with each animal receiving a score from 0-4. 0 meant that there was no labeling; 1 meant that up to 5 cells were labeled per section around the putative injection site; 2 meant that 6-10 cells were labeled per section; 3 meant that 11-30 cells were labeled per section; and 4 meant that over 30 cells were labeled per section.

### Intrapallial injections in Pleurodeles adults

Adult (post-metamorphic) salamanders were deeply anesthetized in 0.1% MS-222. Heads were stabilized throughout surgery and viral injection using a stereotaxic device equipped with custom-built soft ear cups (Kopf Instruments), and animals were kept moist using 0.1% MS-222 soaked gauze. Craniotomies were performed over a region of interest using a dental drill (Foredom) equipped with a 0.75 mm carbide drill bit (Stoelting), and meninges were removed using fine forceps. FastGreen was added to AAV aliquots (as provided by AddGene or NeuroTools) at a ratio of 1:5 for visualization of the injection site. Glass capillary needles were pulled and then broken open using fine forceps (needle tip diameter ranging 5-15 um), backfilled with mineral oil, and then connected to a Nanoject III injection system (Drummond) 100 nL of AAV solution was then pressured injected into neural tissue at an injection rate of 1 nL/12 sec, for a total injection time of approx. 20 min. After injections were complete, the needle was left in the tissue for 3-5 min to prevent leakage, after which the craniotomy was covered with a flexible polymer (PDMS) coverslip, and the skin flap was replaced and sealed with KwikSil (World Precision Instruments). Tissue was perfused and harvested 21-25 days after injection, and fixed overnight at 4 °C in 4% PFA/PBS. 70 μm floating sections were then cut on a vibratome, and stained using immunohistochemistry.

### EdU injections in *Xenopus* tadpoles

EdU administration was performed after intracerebroventricular and/or intraspinoventricular injections of AAV (described above). Anesthetized tadpoles we placed on a moist gauze with the ventral side facing up. A beveled needle was introduced into the peritoneal cavity, and EdU solution was injected (50 mg/kg of body weight). The quantification of overlap between EdU and AAV signal was performed in Imaris on 3D image stacks, each 40 μm thick. The counts were entered into Prism where the plots were generated and statistics calculated as described in Figure legends.

### EdU injections in *Pleurodeles* larvae

Deeply anesthetized larvae were placed on their back in a Sylgard mold after intracerebroventricular injection of the AAV virus. EdU was administered intraperitoneally (50 mg/kg of total body weight), using a glass capillary needle and pressure injection.

### Staining protocols

#### Immunohistochemistry on frozen Xenopus sections

Frozen sections were mounted on glass slides and allowed to dry overnight at 4 °C. The tissue was then rehydrated with 1xPBS+0.2%Triton (PBST) for 2-5 minutes. Following another PBST wash, the tissue was left to incubate with chicken anti-GFP (1:500, Abcam ab13970) and mouse anti-HuC/D (Elavl3/4) (1:500, Invitrogen A21271) in PBST overnight at 4 °C in a humidified chamber. On the following day, sections were moved to room temperature, washed 3 x PBST, and allowed to incubate in Donkey Anti-Chicken IgY conjugated to Alexa 488 (703-545-155, Jackson ImmunoResearch) and Donkey Anti-Mouse IgG conjugated to Cy5 (1:500; 715-175-151, Jackson ImmunoResearch) with DAPI (1:500) in PBST for 45 minutes at RT. Sections were then washed 3 x PBST and once in 1xPBS to remove unbound antibody. EdU staining was performed on sections of the tadpole that received an EdU pulse, by incubating the sections in the Click-iT reaction cocktail for 1 hour according to manufacturers’ instructions (C10340, Invitrogen). Slides were mounted in PVA DABCO antifade medium (10981, Sigma) and left to cure covered for 1 hour at RT. Images were acquired at either a confocal microscope (Zeiss LSM800) or a spinning disk (Nikon CSU W1) using a 20x objective, and processed in Fiji. For size standardization and alignment of images, some images were cropped and placed on a black background in Figure Panels. The quantification of overlap between AAV and Elavl3/4 signal was performed in Image J or Imaris on 3D image stacks, each 40 μm thick. The counts were entered into Prism where the plots were generated and statistics calculated as described in Figure legends.

### Immunohistochemistry on frozen and floating Pleurodeles sections

Frozen sections on glass slides were placed in a humidified chamber, while floating sections were placed in 1 mL of solution in 12-well plates. All sections were blocked at RT for 30 mins - 1 hr in blocking buffer (2.5% BSA, 2.5% sheep serum, 50 mM glycine in PBST (PBS with 0.2% Triton)), and then incubated with chicken anti-GFP (1:500, abcam ab13970), and/or rabbit anti-RFP (1:5000, rockland 600-401-379) in primary Ab solution (10 mM glycine, 0.1% H2O2 in PBST) overnight at 4 °C. Sections were washed 5 x 15 min in PBST at RT, then incubated in Donkey Anti-Chicken IgY conjugated to Alexa 488 (1:500, Jackson ImmunoResearch) and/or Goat Anti-Rabbit conjugated to Alexa 594 (1:500, Fisher) with DAPI (1:1000) in PBST for 2 hrs at RT. Sections were washed 5 x 15 min in PBST at RT and slides were mounted in DAKO fluorescent mounting medium (Agilent Technologies). EdU staining was performed on sections from larvae that received an EdU pulse, after immunohistochemistry, by incubating the sections in the Click-iT reaction cocktail for 45 mins according to manufacturers’ instructions (Invitrogen C10340). Images were acquired using a confocal microscope (Zeiss LSM800) and processed in Fiji. For size standardization and alignment of images, some images were cropped and placed on a black background in Figure Panels.

### Immunohistochemistry on frozen Pelophylax bedriagae sections

Frozen sections were mounted on glass slides and allowed to dry overnight at 4 °C. The tissue was then rehydrated with 1xPBS+0.5%Triton (PBST) for 5 minutes. Following another 3 PBST washes, all sections were blocked at RT for 1 hr in blocking buffer (10% Normal Goat Serum (NGS, ab7481) in 0.3% PBST), and then incubated at 37 °C with Anti-GFP (1:1000, ab290) in blocking solution. Sections were washed 5 x 5 min in 0.1% PBST at RT, then incubated Goat Anti-Rabbit IgG H&L (Alexa Fluor 488)(1:500, ab150077) for 45 min at 37 °C. Following another 3 x 5 min 0.1% PBST washes, the slices were carefully dried and covered with Fluoromount-G with DAPI (Invitrogen Cat. Num 00-4959-52) and covered by coverslip. Images were acquired using a confocal microscope (3i Marianas Spinning Disc) and processed in Fiji. For size standardization and alignment of images, some images were cropped and placed on a black background in Figure Panels.

### Scoring AAV expression in larval screening

For serotype screening in *Pleurodeles* larvae, every 5th section was imaged, acquiring 3 images 1μm apart and a stack was generated in Fiji. Images through the telencephalon and diencephalon were scored as such: low (0-5 cells per section), moderate (6-25 cells per section), and high (more than 25 cells per section).

For serotype screening in Xenopus tadpoles, every 6th section was imaged, acquiring 30-40 images 1.5 μm apart and a maximum intensity projection was generated in Fiji. Brain images were scored as such: low (0-10 cells per section), moderate (11-20 cells per section), and high (more than 20 cells per section).

For serotype screening in *Pelophylax* larvae/tadpoles, each section was stacked acquiring 20-40 images with a 1 μm gap between each image and processed in Fiji. Brain images were scored as such: low (0-5 cells per section), moderate (6-10 cells per section), and high (more than 10 cells per section).

For all positive samples, values 1, 2 and 3 were then assigned to categories low, moderate, and high, respectively, and the average was then calculated for all positive samples.

### Whole mount Hybridization Chain Reaction (HCR) in situ hybridization

HCR-3.0-style probe pairs against the viral DNA/RNA were designed using the insitu_probe_generator ^149^ and ordered from IDT (45 pairs). Brains stored at −20 °C in methanol were equilibrated to 4 °C and bleached overnight in 5% H_2_O_2_ in methanol at 4 °C. The next day, brains were washed in methanol 2 hrs at RT, incubated two times in HCR wash buffer (Molecular Instruments) until the samples sank, and pre-hybridized in hybridization buffer (Molecular Instruments) for 1 hr at 37 °C. Afterwards, brains were incubated 3x overnight at 37 °C in probe hybridization buffer with 6 pmol of the probe set. Excess probe was removed by 3 x 45 min washes in probe wash buffer at 37 °C, and 3 x 45 min washes in 5x SSCT at RT. In case of EdU detection, brains were incubated in the Click-iT reaction cocktail for 45 mins according to manufacturers’ instructions (Invitrogen C10340) at this step, and washed in 5x SSCT for 10 min. The brains were then preamplified for 2 hrs at 4 °C in amplification buffer (Molecular Instruments), and incubated 3x overnight in amplification buffer with snap-cooled hairpins (60 pM, Molecular Instruments) at 4 °C. Primary antibody was also added during amplification when necessary. Excess hairpin was removed by 2x 1 hr washes in 5x SSCT and 2x and overnight washes in 500 mM Tris-HCl pH7.0. Secondary antibody is added during the 5x SSCT washes when necessary. The next day, brains were embedded in 4% agarose in Tris-HCl, and 70 μm vibratome slices were made. Sections were then incubated in DAPI in Tris-HCl for 30 min and mounted in Fluoromount-G® Mounting Medium (SouthernBiotech). Images were acquired using a confocal microscope (Zeiss LSM800) and processed in Fiji.

### Analysis of variable injection outcomes in Pleurodeles adults using factor analysis for mixed data (FAMD)

To analyze injection outcomes as a function of measurable variables in an expanded *Pleurodeles* adult injection dataset, injection metadata were compiled for experiments that were not restricted to the AAV-GFP construct that was used for screening. This expanded dataset included injections of the following AAV constructs: AAV-PHP.eB-CAG-tdTomato (Addgene #59462-PHPeB), AAV-PHP.eB-CAG-GCaMP8s (Neurotools custom virus), AAV-PHP.eB-CAG-hM4D-mCherry (Neurotools custom virus), AAV-PHP.eB-CAG-EGFP (Addgene #37825-PHPeB), AAV-PHP.eB-mDlx-NLS-mRuby2 (Neurotools custom virus), AAVrg-CAG-tdTomato (Addgene #59462-AAVrg), AAVrg-hSyn-EGFP (Addgene #50465-AAVrg), AAVrg-CAG-EGFP (Addgene #37825-AAVrg), AAV9-CAG-EGFP (Addgene #37825-AAV9), and AAV9-hSyn-EGFP (Addgene #50465-AAV9).

Injections of each of these constructs were performed by the same experimenter, using the same injection and floating section immunostaining protocol as described for the serotype screen above. In addition to the weight and age of the animal, injection site, and viral genomes injected, an expression score was assigned to each injected construct. In the case where two constructs were injected in a combined injection solution (dual injection), the outcome for each construct was scored separately (see **Table S3** for full dataset).

Injection outcome scores were assigned on a scale from 0-4, based on individual single-plane confocal images of the putative injection site (i.e. where expression was highest). These scores differed from larval injection outcomes described above, due to the higher-intensity, localized labeling induced by direct intraparenchymal injection. Both qualitative and quantitative data were included in score assignment: A score of 0 was assigned if no labeling was observed, 1 was assigned where <10 cells were seen at the injection site, and labeling intensity was low, 2 was assigned where 10-30 cells were seen, and labeling intensity was intermediate, 3 was assigned where 20-50 cells were seen and labeling intensity was high, and 4 was assigned where >50 cells were seen and labeling intensity was high.

To determine differential contributions of injection variables and outcomes, FAMD analysis and plot visualization was performed in RStudio (R 4.3.2) using the FactoMineR software package ^97^.

