## Supplemental Info for "Adeno-Associated Viral Tools to Trace Neural Development and Connectivity Across Amphibians"

### Supplementary Information

#### Supplementary Discussion

In developing frogs and salamanders, AAV injection into the cerebral ventricle results in GFP expression in neurons three weeks later. As represented in **Figure 3A**, this may occur through transduction of two cell types, radial glia progenitors ("scenario 1") or neurons that were already post-mitotic at the time of injection ("scenario 2").

Taken together, in **developing *Pleurodeles* salamanders injected with AAV-PHP.eB**, our results indicate that <u>both</u> cell types are transduced.

Transduction of radial glia is indicated by the following:

- The presence of AAV DNA/RNA signal in radial glia cells three days after AAV injection (**Figure 3H**) indicates that viral particles were internalized by radial glia cells (1).
- The presence of GFP+ EdU+ neurons three weeks after simultaneous injection of AAV and EdU indicates that GFP+ EdU+ neurons descended from neural progenitors that were still dividing at the time of injection (**Figures 3B-E**).

Transduction of post-mitotic neurons is indicated by the following:

- The presence of AAV DNA/RNA signal in the mantle zone three days after AAV injection (**Figure 3H**) indicates that viral particles were internalized by cells that were either post-mitotic neurons, or neural progenitors in the G2/M phase of their last cell cycle at the time of AAV injection (because mantle zone cells are EdU negative). In the latter case, however, we would expect to see AAV DNA/RNA only in the neurons closest to the ventricular zone. Instead, AAV DNA/RNA signal is present in the entire mantle zone, including the most superficial neurons, which according to our EdU birthdating experiments were born before AAV injection in the experiment in **Figure 3H** (data not shown).
- If AAVs were to transduce only radial glia, in **Figure 3B-C** we would expect to see EdU signal in ~57% of GFP+ neurons. This estimate comes from measurements of cell cycle length in the developing pallium of *Pleurodeles* (2): in early active larvae, the cell cycle length of radial glia is ~10.5 hours, and the S phase is ~6 hours. Therefore radial glia cells spend ~57% of their time in the S phase, when they can incorporate EdU. However, the percentage of GFP+ EdU+ neurons we measured after AAV injection is much lower: ~13% on average, quantified in the pallium for viral injections at stages 41 and 51 (see **Figure 3B-E**; n=1 for each stage, 6 sections each, both hemispheres for the stage 41 injection and the injected hemisphere for the stage 51 injection).

Widespread AAV transduction of post-mitotic neurons is observed in neonatal mice as well, when intracerebroventricular injections are performed at P0, a stage where neurogenesis is long completed. It is believed that the immaturity of the ependymal layer at P0 facilitates the spread of viral particles in the brain parenchyma (3–6). We speculate that a similar mechanism might be in place in developing and post-metamorphic *Pleurodeles*.

However, in **developing *Xenopus* frogs injected with AAV5**, our results suggest the mechanism of isochronic cohorting may differ. Our data support scenario 1: the hypothesis that labeling of neurons three weeks after AAV injection is primarily the result of AAV transduction of radial glia progenitors. Unlike in *Pleurodeles*, in *Xenopus* we observe AAV-driven GFP expression in radial glia lining the ventricular zone in both the brain and spinal cord (**Figure 2B', G'; Figure S2I-J**). In addition, if AAVs were to only transduce radial glia, one would expect ~20% of GFP+EdU+ neurons after AAV-GFP/EdU co-injection. This calculation derives from the observed *Xenopus* CNS cell cycle duration of 35-40 hours and S-phase duration of 5-7 hours at NF stage 50-54 (7). Remarkably, this predicted overlap maps almost exactly with what we observe experimentally at this same stage, as ~25% of AAV-driven GFP+ cells are also EdU+ across the CNS (**Figure S2B**).

The difference in radial glia transduction and EdU/AAV overlap in frogs and salamanders raise the interesting question of why AAV viruses would behave differently in the two species. One potential reason for this might be the difference between the AAV serotypes - with AAV5 and AAV-PHP.eB being the most evolutionarily and mechanistically distinct viruses. AAV5, for example, uses a different receptor combination for cell entry as compared to AAV-PHP.eB (see **Discussion**). These differences could also be further propagated by species-specific variations in either organismal cellular features or viral expression.

Taking all data in salamander and frog together, **both AAV-PHP.eB and AAV5 target temporal cohorts upon injection into the ventricle**, however, this targeting may occur through distinct mechanisms in the two species.


1. Dhungel BP, Bailey CG, Rasko JEJ. Journey to the center of the cell: tracing the path of AAV transduction. Trends Mol Med. 2021 Feb;27(2):172–84.
2. Joven A, Wang H, Pinheiro T, Hameed LS, Belnoue L, Simon A. Cellular basis of brain maturation and acquisition of complex behaviors in salamanders. Development. 2018 Jan 8;145(1).
3. Passini MA, Wolfe JH. Widespread gene delivery and structure-specific patterns of expression in the brain after intraventricular injections of neonatal mice with an adeno-associated virus vector. J Virol. 2001 Dec;75(24):12382–92.
4. Chakrabarty P, Rosario A, Cruz P, Siemienski Z, Ceballos-Diaz C, Crosby K, et al. Capsid serotype and timing of injection determines AAV transduction in the neonatal mice brain. PLoS ONE. 2013 Jun 25;8(6):e67680.
5. Passini MA, Watson DJ, Vite CH, Landsburg DJ, Feigenbaum AL, Wolfe JH. Intraventricular brain injection of adeno-associated virus type 1 (AAV1) in neonatal mice results in complementary patterns of neuronal transduction to AAV2 and total long-term correction of storage lesions in the brains of beta-glucuronidase-deficient mice. J Virol.

2003 Jun;77(12):7034–40.

6. Kim J-Y, Grunke SD, Levites Y, Golde TE, Jankowsky JL. Intracerebroventricular viral injection of the neonatal mouse brain for persistent and widespread neuronal transduction. J Vis Exp. 2014 Sep 15;(91):51863.

7. Thuret R, Auger H, Papalopulu N. Analysis of neural progenitors from embryogenesis to juvenile adult in Xenopus laevis reveals biphasic neurogenesis and continuous lengthening of the cell cycle. Biol Open. 2015 Nov 30;4(12):1772–81.

**Figure S1. Further AAV screening in larval amphibian brains and cell culture.**
**A-I.** Analysis of AAV.PHP.eB-CAG-GFP transduction efficiency after intracerebroventricular (ICV) injection in the developing *Pleurodeles* brain. **A**. Experimental design: ICV injection of PHP.eB-CAG-GFP and intraperitoneal (IP) injection of EdU in an early-active larva (stage 41), and analysis 23 days later. **B-I**. Coronal sections through the *Pleurodeles* brain showing GFP (green) and EdU (magenta) labeled cells in the fore- and midbrain. Scale bars are 200 um. DAPI, blue. **J-M.** CAG/Syn expression comparison and confirmation of neuronal identity with cell type marker, Elavl3/4, in *Xenopus.* **J.** Schematic overview of the experimental setup: ICV injection of AAV5-CAG-GFP or AAV5-hSyn-GFP in metamorphic tadpoles, and analysis 21 days later. Coronal sections through the telencephalon showing the labeling pattern of AAV5 with a CAG (**K**) and hSyn promoter (**L**). Magnifications show the overlap between the AAV labeling and a neuronal marker, Elavl3/4. AAV-driven GFP, green. Elavl3/4, magenta. **M.** Quantification shows the mean ± standard deviation (SD) by animal of the percentage of Elavl3/4 and AAV double-positive cells, relative to the total number of AAV-positive cells, for all conditions tested. Each dot represents the average number or percentage of cells in a 40 µm thick section calculated from >3 full-stack images; n = 2 animals. **N-Q.** AAVs transduce cells in late embryonic stage *Xenopus* brain. **N**. Schematic overview of the experimental design: ICV injections of AAVrg-hSyn-GFP, AAV5-hSyn-GFP, or AAV1-CAG-GFP into NF40-41 *Xenopus* larvae, and analysis 21 days later. **O-Q.** Coronal sections through the *Xenopus* brain showing GFP labeled cells in the fore- and midbrain, section levels shown in **N**. Scale bars represent 200 um. AAV-driven GFP, green. DAPI, blue. **R.** *In vitro* screening of AAV serotypes: AAV1-CAG-GFP (green) efficiently transduces a frog epithelial cell line, XLJ-1, after 5 days of incubation at the concentration of 0.0125 µL virus/µL media.

**Abbreviations**: A, anterior; d, day; ICV, intracerebroventricular injection; IP, intraperitoneal injection; ns, not significant; P, posterior; St, stage; TL, transmitted light.

**Figure S2. AAV labels different populations of neurons or radial glia based on the developmental stage at the time of intraventricular injection. A.** Schematic overview of the experimental design. Prometamorphic *Xenopus laevis* tadpoles NF stage 49-50 were co-injected with AAV5-CAG-GFP intracerebroventricularly (ICV) and intraspinoventricularly (ISV), and with EdU intraperitoneally (IP). Tissue was analyzed three weeks later, GFP and EdU signals were imaged, and the percentage of AAV+ EdU+ cells was quantified. **B.** Quantification of AAV and EdU overlap shows the mean percentage ± standard deviation (SD) by animal of AAV and EdU double positive cells, relative to total number of AAV-positive cells, for all CNS or a specific region. Each dot represents the average percentage of cells in a 40 µm thick section calculated from >2 full-stack images; n = 2-3 animals. **C-D**. Representative coronal sections are shown for different regions of the *Xenopus* brain with section planes indicated on the schematic in **A**. Arrowsheads in the insets **C'** and **D'** indicate cells with overlapping GFP, EdU and Elavl3 (neuronal marker) signal at a given optical plane. **E–L.** Prometamorphic tadpoles were injected with AAV5-CAG-GFP intraspinoventricularly at either NF52 (**E-F**) or NF57 (**G-L**) and GFP expression was analyzed 3 weeks later. The injection at the earlier stage of metamorphosis labels both ventral and dorsal interneurons (**F**), whereas the later injection marks the dorsal population (**H**), which is in line with the known birth order of spinal cell types. In case of a developmentally delayed animal that remained at NF57 for three weeks (**I**), a growth stall that is frequently observed in *Xenopus* development, labeling is largely confined to the radial glia (**J**), suggesting that AAV injected into the spinal canal labels the proliferating progenitors at the time of injection. In contrast, intraparenchymal injection of the spinal cord (**K**) labeled the postmitotic LMC and MMC motor neurons (**L**), identified by their characteristic morphology and columnar position, in addition to dorsal interneurons. Scale bars in brain overview images and magnifications represent 400 µm and 50 µm, respectively. All sections are 40 µm thick and cut at a coronal (brain) or horizontal (spinal cord) plane. AAV-driven GFP, green. EdU, magenta. Elavl3/4, cyan. DAPI, blue.

**Abbreviations:** A, anterior; BR, brachial; d, day; Di/Dien, diencephalon; Hyp, hypothalamus; ICV, intracerebroventricular injection; IP, intraperitoneal injection; ISV, intraspinoventricular injection; LMC, lateral motor column; LU, lumbar; Me/Mes, mesencephalon; ML, mitral layer; MMC, medial motor column; P, posterior; PL, progenitor layer; Rh, rhombencephalon; SC, spinal cord; Tel, telencephalon; Th, thalamus.

**Figure S3. Intracerebroventricular injections produce widespread AAV expression in post-metamorphic *Pleurodeles.* A.** Schematic showing intracerebroventricular (ICV) injection of AAV-PHP.eB-CAG-GFP into the left telencephalic hemisphere in a 3 month-old post-metamorphic *Pleurodeles*. **B-H.** Representative coronal sections imaged along the anterior-posterior axis (section levels shown in **A**), with insets from each hemisphere shown below. **I-J.** Widespread neuronal labeling after ICV AAV injection of AAV-PHP.eB-CAG-GFP, with inset showing sporadic labeling in ependymoglial cells (EGCs) lining the ventricular zone (VZ). **K-L.** Electroporation of CAG-GFP plasmid produces effective labeling of EGCs, co-stained with Sox2 (red). AAV-driven GFP, green. Sox2, magenta. DAPI, blue. Scale bars in overview images and magnifications are 200 µm and 100 µm, respectively.

**Abbreviations:** A, anterior; D, dorsal; Di, diencephalon; Me, mesencephalon; P, posterior; Rh, rhombencephalon; Tel, telencephalon; V, ventral; VZ, ventricular zone.

**Figure S4. Sources of variability in *Pleurodeles* adult injection outcomes. A.** Adult *Pleurodeles* were injected bilaterally into dorsal pallium with AAV9-CAG-GFP (left hemisphere) and AAV9-hSyn-GFP (right hemisphere), to compare expression strength as a function of promoter choice. Single coronal section showing expression of both constructs. **B-C.** Insets showing expression of AAV9-CAG-GFP (**B**) and AAV9-hSyn-GFP (**C**), imaged with a confocal microscope under the same settings. Despite differences in strength, both constructs drove detectable expression of GFP. Scale bar is 200 µm in B, and 100 µm in C-D. **D-E.** Adult *Pleurodeles* were injected into dorsal pallium with AAV9-CAG-GFP, and then adjacent sections (70 µm apart) were separated to test the necessity of GFP immunohistochemistry (IHC) for detecting reporter fluorescence. IHC for GFP yielded high-intensity labeling (**D**), whereas imaging in the absence of IHC using the same confocal settings yielded detectable expression, but lower fluorescence intensity. In D-E, scale bars are 200 µm; AAV-driven GFP, green. DAPI, blue. **F.** Screeplot showing the percentage (%) of explained variance for each dimension following factor analysis of mixed data (FAMD) of metadata for 25 animals and 41 injection outcomes. FAMD included continuous variables (total v.g., age, weight, and outcome score) and categorical variables (reporter, manufacturer, promoter, serotype, single vs. dual injection, and injection site). **G.** Contribution of each variable to the first FAMD dimension (Dim. 1). Reporter, manufacturer, total viral genomes (v.g.), and age contributed above chance (outlined in red, dotted red line >10%). **H.** Contribution of each variable to the second FAMD dimension (Dim. 2). Promoter, reporter, expression score and serotype contributed above chance (outlined in red, dotted red line >10%). **I-R.** Plots showing each individual injection in FAMD space. Each data point is color-coded according to one of the quantitative or qualitative variables included in the FAMD analysis. **S.** Linear regression of expression score against age, showing a significant negative correlation between animal age and quality of injection outcomes (score); $F(1,40) = 9.81$, $p = 0.0032$, $R^2=0.18$.

**Abbreviations:** AG, AddGene; DP, dorsal pallium; GC8s, GCaMP8s; IHC, immunohistochemistry; mCh, hM4D-mCherry; mR2, NLS-mRuby2; NT, Neurotools; OB, olfactory bulb; tdT, tdTomato; Th, Thalamus; v.g., viral genomes; Wgt: weight.

**Figure S5. Retrograde tracing of intrapallial connectivity in *Xenopus laevis*. A.** Post-metamorphic *Xenopus* was injected into the caudal dorsal pallium with AAVrg-CAG-GFP (right hemisphere), and the spread of labeling was evaluated after GFP immunostaining. Representative coronal sections of the telencephalon imaged along the anterior-posterior axis are shown in **B-H.**, with insets highlighting each hemisphere shown below. Robust cell body and axonal labeling is present in the ipsilateral DP injection site with traced cell bodies found in both ipsilateral and contralateral MP, DP, LP, and septum. The distance between the most anterior (**B**) and the most posterior (**H**) labeled neurons was approximately 2 mm. Shown are maximum intensity projections of 40 µm thick cryosections. Scale bars in overview images and magnifications represent 400 and 50 µm, respectively. AAV-driven GFP, green. DAPI, blue.

**Abbreviations:** A, anterior; APOA, anterior preoptic area; CeA, central amygdala; D, dorsal; Di, diencephalon; DP, dorsal pallium; LA, lateral amygdala; LP, lateral pallium; Me, mesencephalon; MP, medial pallium; P, posterior; Sept, septum; Tel, telencephalon; V, ventral; VP, ventral pallium.

**Table S1.** Overview of cross-species AAV screen performed in *Pelophylax bedrigae, Xenopus laevis,* and *Pleurodeles waltl.* Injections and scoring (none, low, moderate, high) were performed for each species according to **STAR Methods**. For each viral construct and species, stage, injection location, sample size, and injection outcome is reported. Percentage of success was calculated as a function of number of samples with labeling to sample size.

**Table S2.** Summary of injections in adult *Xenopus laevis.* Shown are the post metamorphic stage, body weight, type of injection, injected virus details, volume, manufacturer, injection site and the expression score (0-4).

**Table S3.** Summary of injections in adult *Pleurodeles waltl* included in factor analysis of mixed data (FAMD). Continuous variables were age (days), weight (g), injected volume (nl), total viral genomes (v.g.), and expression score (0-4). Categorical variables were single vs. dual constructs injected, serotype, promoter, reporter, manufacturer, and injection site.

**Table S4.** HCR probe sets

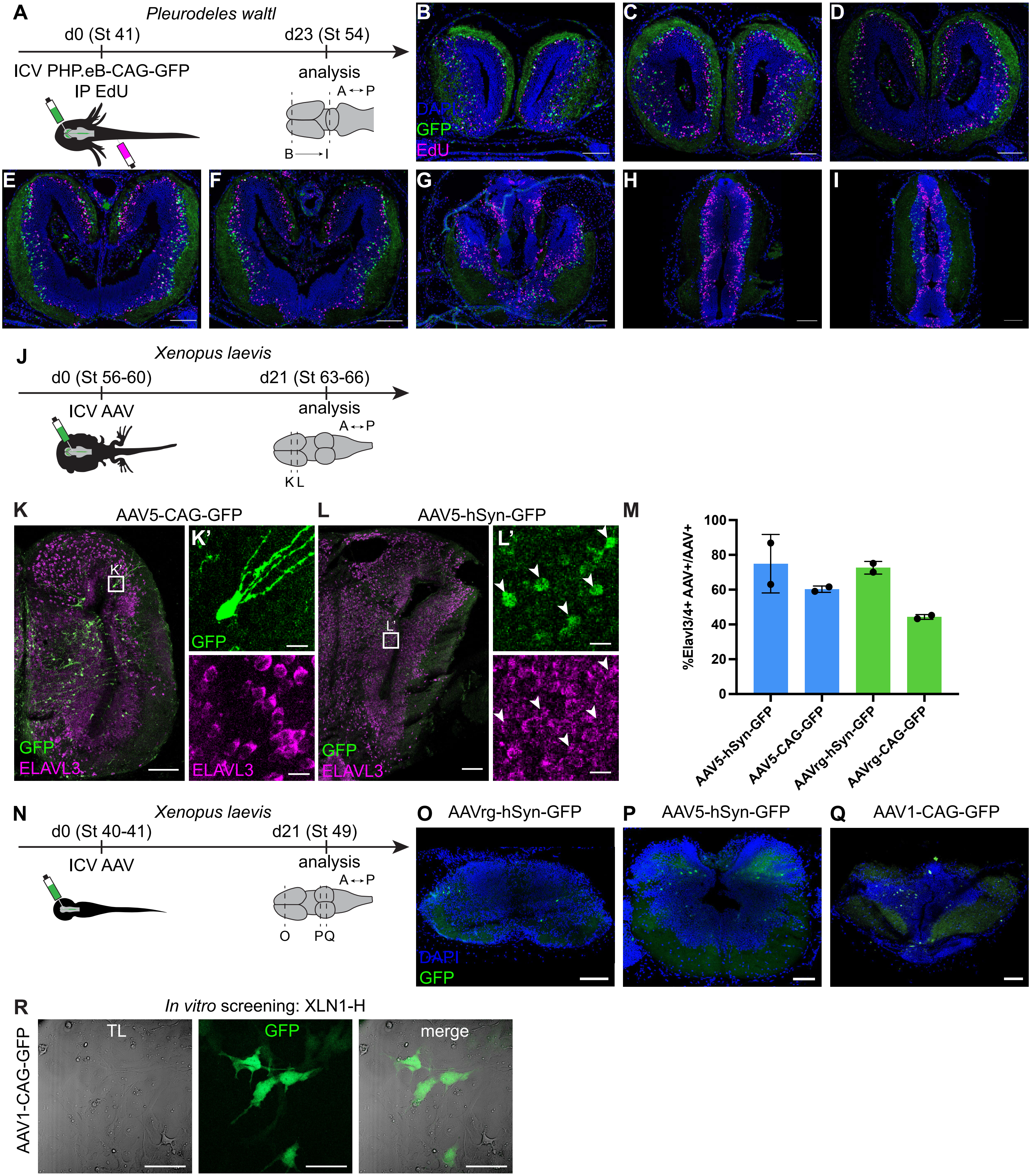


**Supplementary Figure S1**

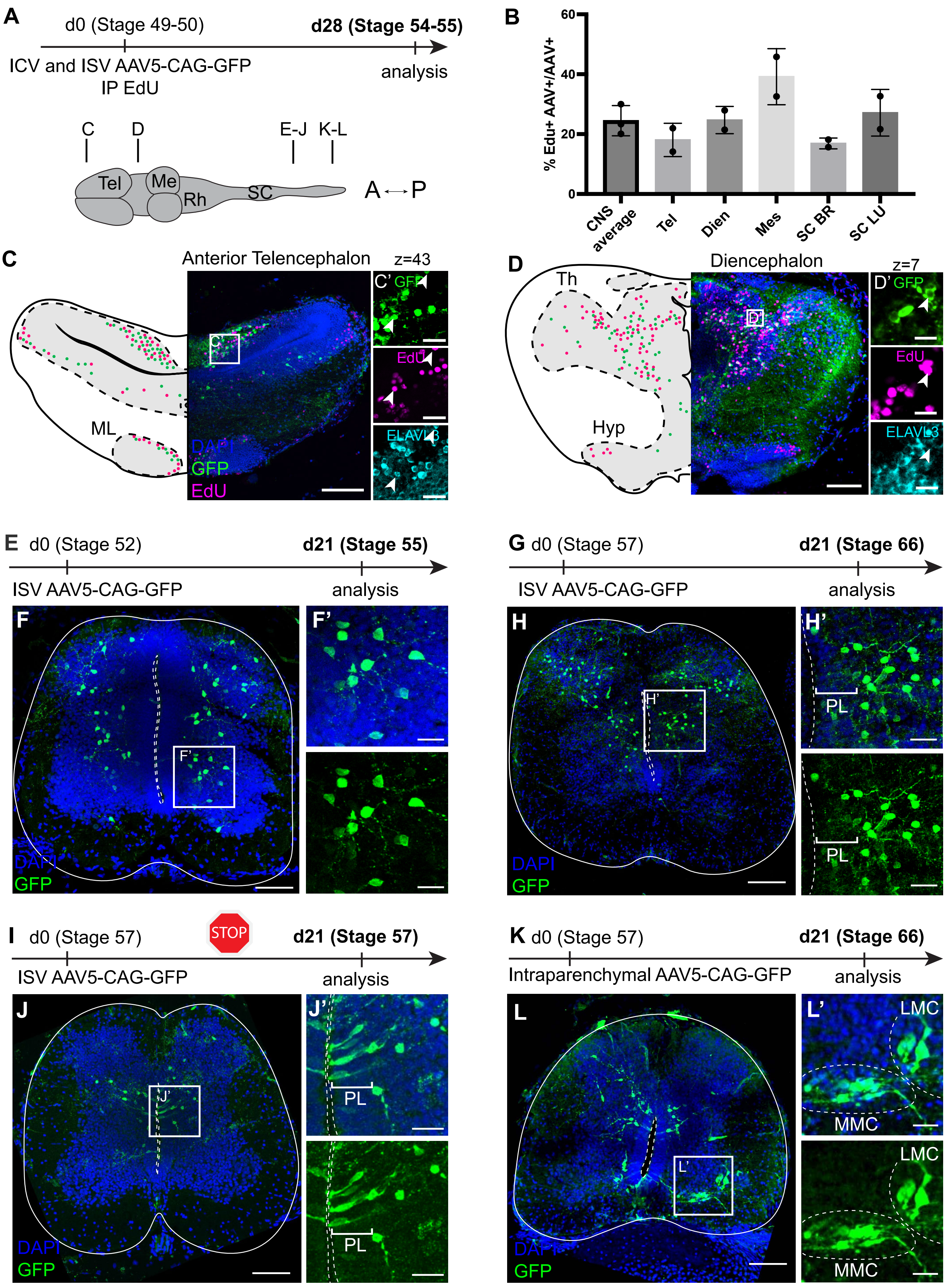


**Supplementary Figure S2**

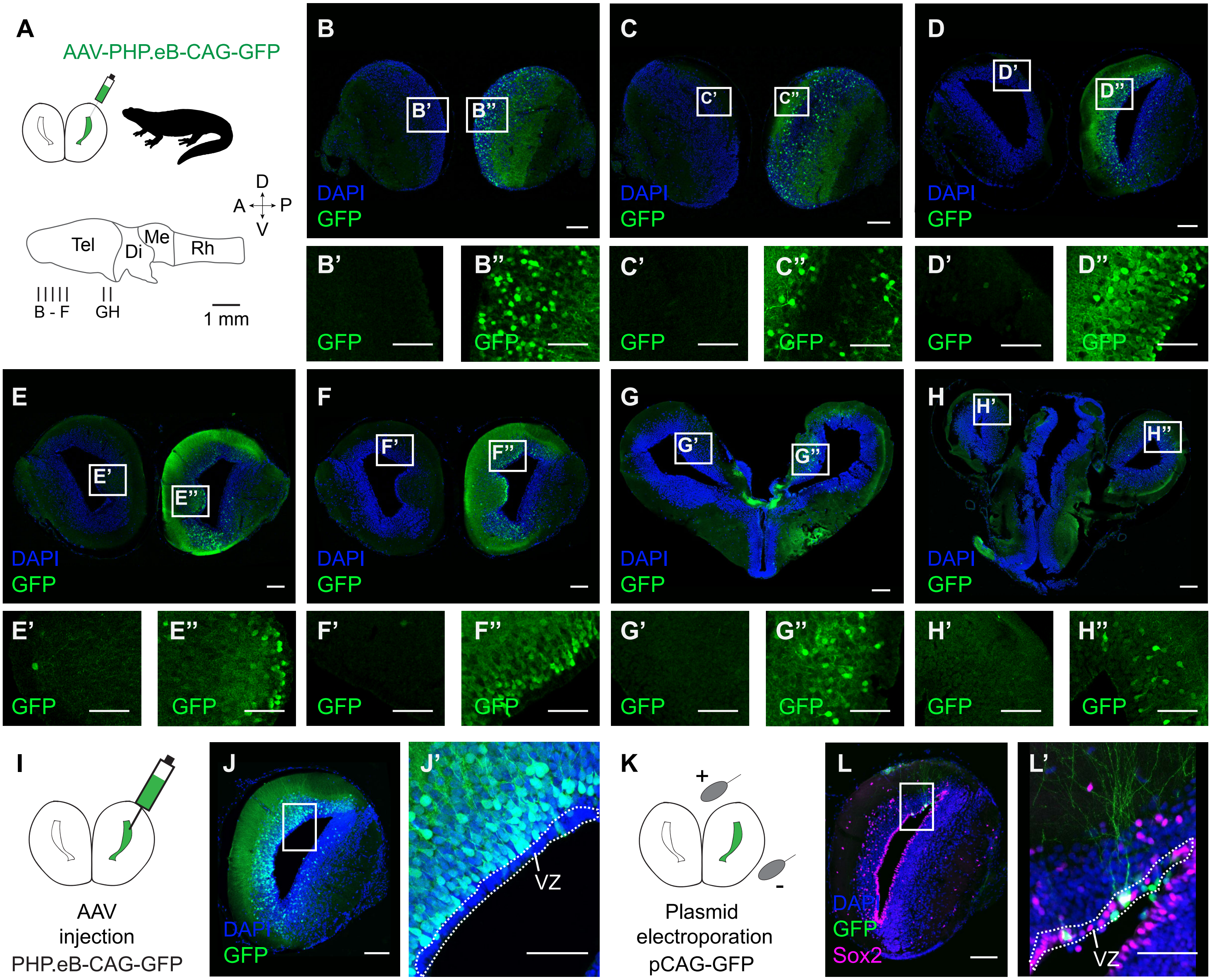


**Supplementary Figure S3**

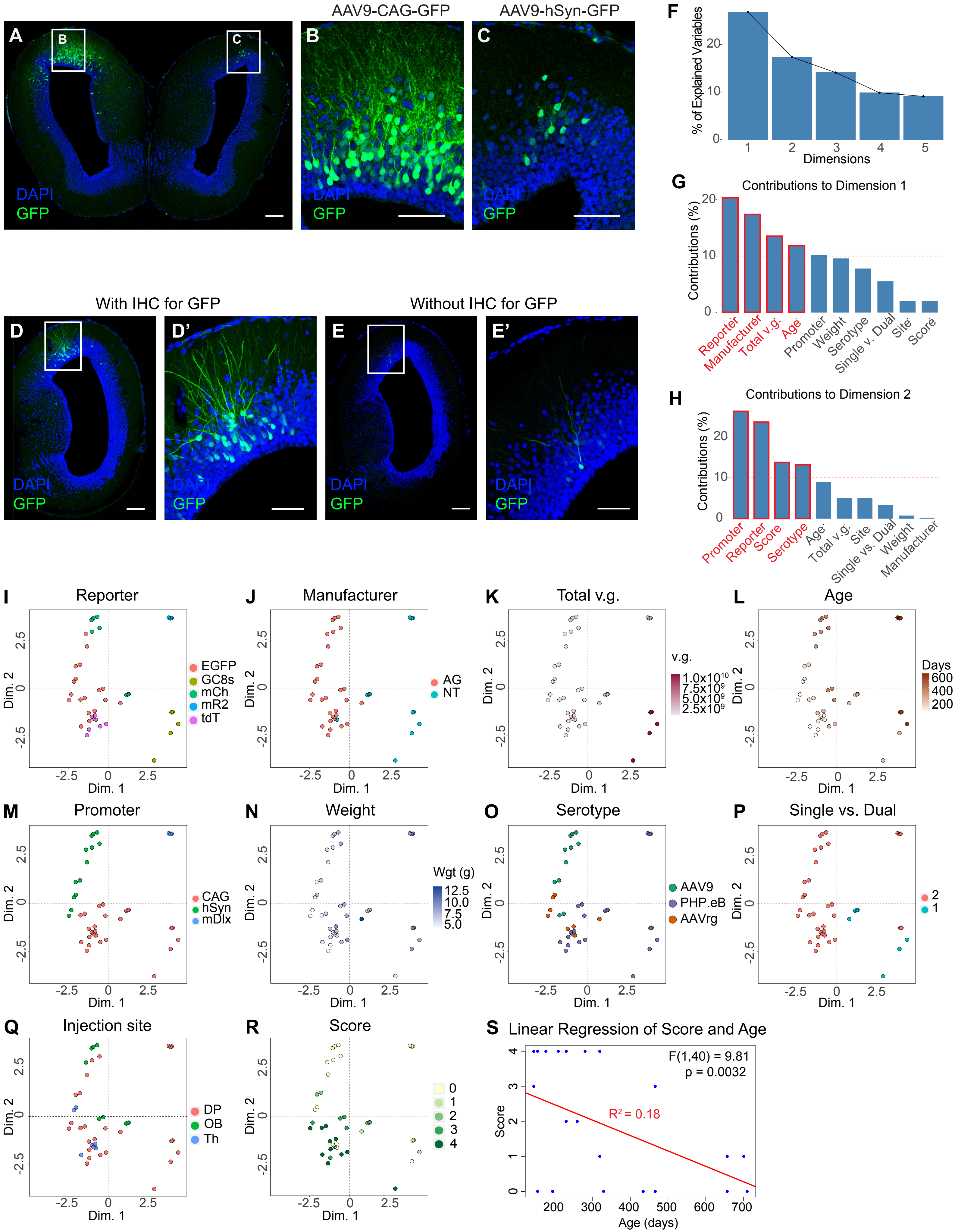


**Supplementary Figure S4**

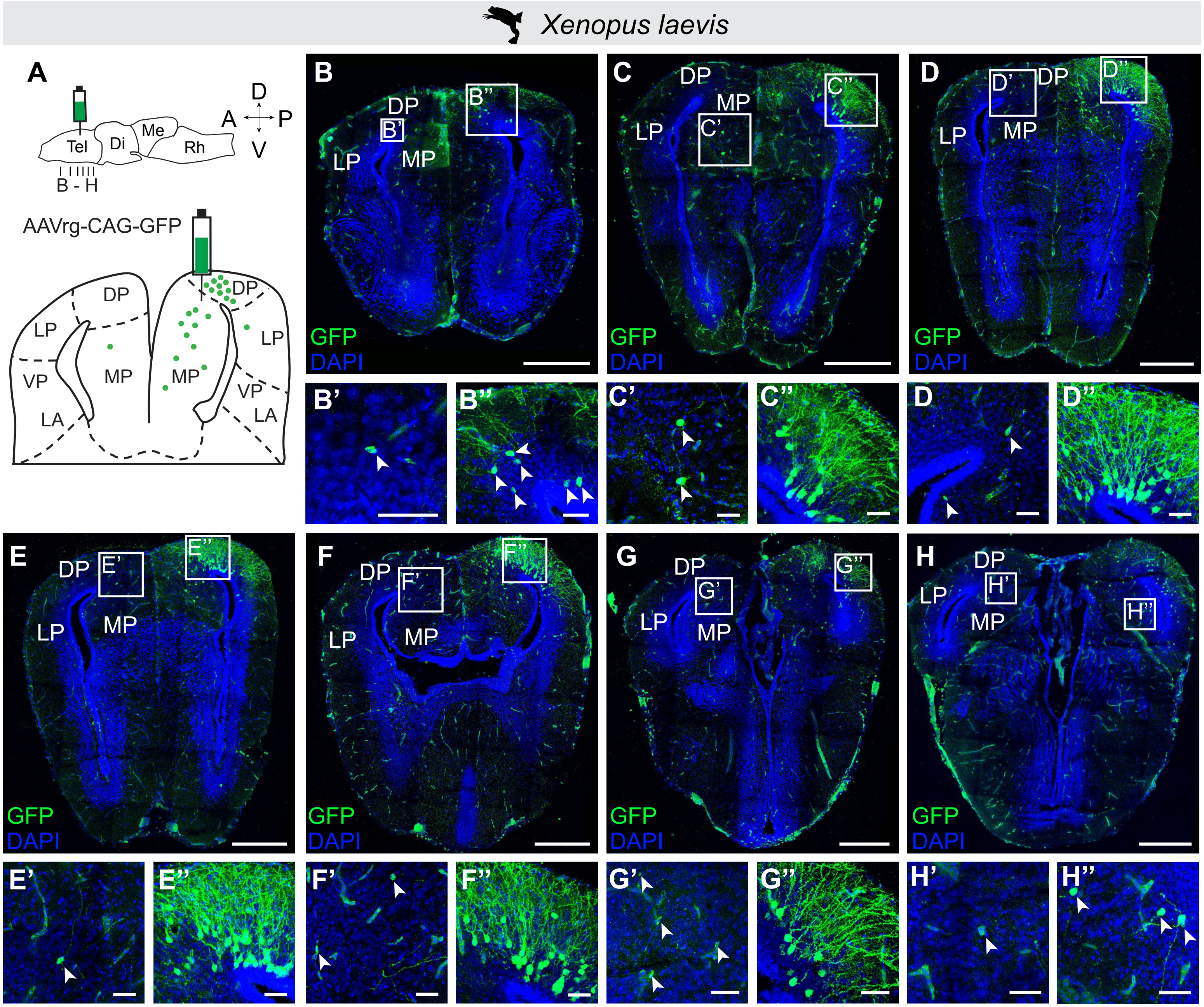


**Supplemental Figure S5**

| Virus | Species | Stage | Injection location | Sample size | # Without Labeling | # With Labeling | % Success | Ratio | # None | # Low | # Moderate | # High |
|---|---|---|---|---|---|---|---|---|---|---|---|---|
| AAV-PHP.eB-CAG-GFP | *Pelophylax bedriagae* | Adult | Brain ventricle | 6 | 6 | 0 | 0 | 0/6 | 6 | 0 | 0 | 0 |
| AAV1-hSyn-GFP | *Pelophylax bedriagae* | Adult | Brain ventricle | 1 | 1 | 0 | 0 | 0/1 | 1 | 0 | 0 | 0 |
| AAV5-CAG-GFP | *Pelophylax bedriagae* | Adult | Brain ventricle | 2 | 2 | 0 | 0 | 0/2 | 2 | 0 | 0 | 0 |
| AAV5-hSyn-GFP | *Pelophylax bedriagae* | Adult | Brain ventricle | 1 | 1 | 0 | 0 | 0/1 | 1 | 0 | 0 | 0 |
| AAV9-hSyn-GFP | *Pelophylax bedriagae* | Adult | Brain ventricle | 1 | 1 | 0 | 0 | 0/1 | 1 | 0 | 0 | 0 |
| AAVrg-hSyn-GFP | *Pelophylax bedriagae* | Adult | Brain ventricle | 5 | 5 | 0 | 0 | 0/5 | 5 | 0 | 0 | 0 |
| AAV-PHP.eB-CAG-GFP | *Pelophylax bedriagae* | Tadpole | Brain ventricle | 7 | 6 | 1 | 14 | 1/7 | 6 | 0 | 1 | 0 |
| AAV5-CAG-GFP | *Pelophylax bedriagae* | Tadpole | Brain ventricle | 7 | 6 | 1 | 14 | 1/7 | 6 | 1 | 0 | 0 |
| AAVrg-hSyn-GFP | *Pelophylax bedriagae* | Tadpole | Brain ventricle | 5 | 5 | 0 | 0 | 0/5 | 5 | 0 | 0 | 0 |
| AAV-PHP.eB-CAG-GFP | *Pleurodeles waltl* | Adult | Pallium | 5 | 0 | 5 | 100 | 5/5 | 0 | 0 | 1 | 4 |
| AAV1-hSyn-GFP | *Pleurodeles waltl* | Adult | Pallium | 2 | 2 | 0 | 0 | 2/2 | 2 | 0 | 0 | 0 |
| AAV5-hSyn-GFP | *Pleurodeles waltl* | Adult | Pallium | 2 | 2 | 0 | 0 | 2/2 | 2 | 0 | 0 | 0 |
| AAV8-hSyn-GFP | *Pleurodeles waltl* | Adult | Pallium | 2 | 2 | 0 | 0 | 2/2 | 2 | 0 | 0 | 0 |
| AAV9-hSyn-GFP | *Pleurodeles waltl* | Adult | Pallium | 6 | 2 | 4 | 67 | 4/6 | 2 | 0 | 2 | 2 |
| AAV9-CAG-GFP | *Pleurodeles waltl* | Adult | Pallium | 2 | 0 | 2 | 100 | 2/2 | 0 | 0 | 0 | 2 |
| AAVrg-hSyn-GFP | *Pleurodeles waltl* | Adult | Pallium | 4 | 0 | 4 | 100 | 4/4 | 0 | 1 | 1 | 2 |
| AAVrg-CAG-GFP | *Pleurodeles waltl* | Adult | Pallium | 4 | 0 | 4 | 100 | 4/4 | 0 | 2 | 0 | 2 |
| AAV-PHP.eB-CAG-GFP | *Pleurodeles waltl* | Tadpole | Brain ventricle | 13 | 0 | 13 | 100 | 13/13 | 0 | 0 | 0 | 13 |
| AAV1-hSyn-GFP | *Pleurodeles waltl* | Tadpole | Brain ventricle | 2 | 0 | 2 | 100 | 2/2 | 0 | 2 | 0 | 0 |
| AAV2-hSyn-GFP | *Pleurodeles waltl* | Tadpole | Brain ventricle | 5 | 0 | 5 | 100 | 5/5 | 0 | 4 | 1 | 0 |
| AAV5-hSyn-GFP | *Pleurodeles waltl* | Tadpole | Brain ventricle | 2 | 0 | 2 | 100 | 2/2 | 0 | 2 | 0 | 0 |
| AAV9-hSyn-GFP | *Pleurodeles waltl* | Tadpole | Brain ventricle | 4 | 0 | 4 | 100 | 4/4 | 0 | 0 | 0 | 4 |
| AAVrg-hSyn-GFP | *Pleurodeles waltl* | Tadpole | Brain ventricle | 3 | 0 | 3 | 100 | 3/3 | 0 | 0 | 3 | 0 |
| AAV5-CAG-GFP | *Xenopus laevis* | Juvenile/Adult | Pallium | 11 | 4 | 7 | 64 | 7/11 | 4 | 0 | 1 | 6 |
| AAVrg-CAG-GFP | *Xenopus laevis* | Juvenile/Adult | Pallium | 3 | 0 | 3 | 100 | 3/3 | 0 | 1 | 0 | 2 |
| AAV-PHP.eB-CAG-GFP | *Xenopus laevis* | Tadpole | CNS ventricle | 8 | 8 | 0 | 0 | 0/8 | 8 | 0 | 0 | 0 |
| AAV1-CAG-GFP | *Xenopus laevis* | Tadpole | CNS ventricle | 4 | 2 | 2 | 50 | 2/4 | 2 | 1 | 1 | 0 |
| AAV1-hSyn-GFP | *Xenopus laevis* | Tadpole | CNS ventricle | 7 | 5 | 2 | 29 | 2/7 | 5 | 2 | 0 | 0 |
| AAV2-hSyn-GFP | *Xenopus laevis* | Tadpole | CNS ventricle | 5 | 5 | 0 | 0 | 0/5 | 5 | 0 | 0 | 0 |
| AAV5-CAG-GFP | *Xenopus laevis* | Tadpole | CNS ventricle | 12 | 1 | 11 | 92 | 11/12 | 1 | 0 | 1 | 10 |
| AAV5-hSyn-GFP | *Xenopus laevis* | Tadpole | CNS ventricle | 6 | 0 | 6 | 100 | 6/6 | 0 | 2 | 0 | 4 |
| AAV8-hSyn-GFP | *Xenopus laevis* | Tadpole | CNS ventricle | 6 | 6 | 0 | 0 | 0/6 | 6 | 0 | 0 | 0 |
| AAV9-hSyn-GFP | *Xenopus laevis* | Tadpole | CNS ventricle | 11 | 11 | 0 | 0 | 0/11 | 11 | 0 | 0 | 0 |
| AAVrg-CAG-GFP | *Xenopus laevis* | Tadpole | CNS ventricle | 2 | 1 | 1 | 50 | 1/2 | 1 | 1 | 0 | 0 |
| AAVrg-hSyn-GFP | *Xenopus laevis* | Tadpole | CNS ventricle | 6 | 2 | 4 | 67 | 4/6 | 2 | 0 | 1 | 3 |

**Table S1.** Overview of cross-species AAV screen performed in *Pelophylax bedrigae, Xenopus laevis,* and *Pleurodeles waltl.* Injections and scoring (none, low, moderate, high) were performed for each species according to **Methods**. For each viral construct and species, stage, injection location, sample size, and injection outcome is reported. % success was calculated as a function of # of samples with labeling to sample size.

| Animal # | Dev. stage | Weight (g) | Single vs dual | Serotype | Promoter | Reporter | Titer(v.g./mL) | Volume (nL) | Manufacturer | Injection site | Expression score |
|---|---|---|---|---|---|---|---|---|---|---|---|
| 1 | PM 0 | 1.79 | single | AAV5 | CAG | EGFP | 1.00E+13 | 100 | Addgene | MP | 3 |
| 2 | PM 0 | 2.47 | single | AAV5 | CAG | EGFP | 1.00E+13 | 100 | Addgene | MP | 0 |
| 3 | PM 1 | 3.25 | single | AAV5 | CAG | EGFP | 1.00E+13 | 100 | Addgene | MP | 0 |
| 4 | PM 0 | 1.19 | single | AAV5 | CAG | EGFP | 1.00E+13 | 100 | Addgene | MP | 0 |
| 5 | PM 0 | 1.39 | single | AAV5 | CAG | EGFP | 1.00E+13 | 100 | Addgene | MP | 3 |
| 6 | PM 0 | 1.08 | single | AAVrg | CAG | EGFP | 2.20E+13 | 100 | Addgene | DP/MP | 4 |
| 7 | PM 0 | 1.24 | single | AAVrg | CAG | EGFP | 2.20E+13 | 100 | Addgene | DP/MP | 4 |
| 8 | PM 0 | 1.36 | single | AAVrg | CAG | EGFP | 2.20E+13 | 100 | Addgene | DP/MP | 1 |
| 9 | PM 2/3 | 12.21 | single | AAV5 | CAG | EGFP | 1.00E+13 | 100 | Addgene | telencephalon | 4 |
| 10 | PM 4/5 | 7.85 | single | AAV5 | CAG | EGFP | 1.00E+13 | 100 | Addgene | telencephalon | 2 |
| 11 | PM 4/5 | 5.72 | single | AAV5 | CAG | EGFP | 1.00E+13 | 100 | Addgene | telencephalon | 3 |
| 12 | PM 4 | 8.69 | single | AAV5 | CAG | EGFP | 1.00E+13 | 100 | Addgene | telencephalon | 1 |
| 13 | PM 5 | 7.4 | single | AAV5 | CAG | EGFP | 1.00E+13 | 100 | Addgene | telencephalon | 3 |
| 14 | PM 3 | 10.83 | single | AAV5 | CAG | EGFP | 1.00E+13 | 100 | Addgene | telencephalon | 0 |

**Table S2.** Summary of injections in adult *Xenopus laevis.* Shown are the post metamorphic stage, body weight, type of injection, injected virus details, volume, manufacturer, injection site and the expression score (0-4).

| Animal ID | Age (days) | Weight (g) | Single vs dual | Serotype | Promoter | Reporter | Titer(v.g./mL) | Volume (nl) | Total v.g. | Manufacturer | Injection site | Expression Score |
|---|---|---|---|---|---|---|---|---|---|---|---|---|
| ej23 | 230 | 5.4 | dual | AAV9 | CAG | EGFP | 2.60E+13 | 100 | 1.30E+09 | Addgene | DP | 4 |
| ej24 | 230 | 7.2 | dual | AAV9 | CAG | EGFP | 2.60E+13 | 100 | 1.30E+09 | Addgene | DP | 4 |
| ej1 | 434 | 6.3 | dual | AAV9 | hSyn | EGFP | 1.90E+13 | 100 | 9.50E+08 | Addgene | DP | 0 |
| ej2 | 434 | 6.9 | dual | AAV9 | hSyn | EGFP | 1.30E+13 | 100 | 6.50E+08 | Addgene | OB | 0 |
| ej23 | 230 | 5.4 | dual | AAV9 | hSyn | EGFP | 1.90E+13 | 100 | 9.50E+08 | Addgene | DP | 2 |
| ej24 | 230 | 7.2 | dual | AAV9 | hSyn | EGFP | 1.90E+13 | 100 | 9.50E+08 | Addgene | DP | 2 |
| ej1 | 434 | 6.3 | dual | AAV9 | hSyn | hM4D-mCherry | 2.30E+13 | 100 | 1.15E+09 | Addgene | DP | 0 |
| ej2 | 434 | 6.9 | dual | AAV9 | hSyn | hM4D-mCherry | 2.30E+13 | 100 | 1.15E+09 | Addgene | OB | 0 |
| ej3 | 466 | 7.2 | dual | AAV9 | hSyn | hM4D-mCherry | 2.30E+13 | 100 | 1.15E+09 | Addgene | OB | 0 |
| ej4 | 466 | 8.9 | dual | AAV9 | hSyn | hM4D-mCherry | 2.30E+13 | 100 | 1.15E+09 | Addgene | OB | 0 |
| ej5 | 466 | 9.1 | dual | AAV9 | hSyn | hM4D-mCherry | 2.30E+13 | 100 | 1.15E+09 | Addgene | DP | 0 |
| ej25 | 319 | 7.74 | dual | AAVrg | CAG | EGFP | 2.00E+13 | 150 | 1.50E+09 | Addgene | DP | 4 |
| ej26 | 319 | 10.25 | dual | AAVrg | CAG | EGFP | 2.00E+13 | 150 | 1.50E+09 | Addgene | DP | 4 |
| ej27 | 319 | 14.92 | single | AAVrg | CAG | EGFP | 2.00E+13 | 100 | 2.00E+09 | Addgene | DP | 1 |
| ej20 | 154 | 6.1 | dual | AAVrg | CAG | tdTomato | 2.60E+13 | 150 | 1.95E+09 | Neurotools | Th | 0 |
| ej18 | 144 | 4.9 | dual | AAVrg | hSyn | EGFP | 2.20E+13 | 100 | 1.10E+09 | Addgene | DP | 4 |
| ej19 | 144 | 5.3 | dual | AAVrg | hSyn | EGFP | 2.20E+13 | 100 | 1.10E+09 | Addgene | DP | 3 |
| ej21 | 194 | 7.8 | dual | AAVrg | hSyn | EGFP | 1.30E+13 | 150 | 9.75E+08 | Addgene | Th | 0 |
| ej22 | 194 | 6.4 | dual | AAVrg | hSyn | EGFP | 2.20E+13 | 150 | 1.65E+09 | Addgene | Th | 0 |
| ej20 | 154 | 6.1 | dual | PHP.eB | CAG | EGFP | 1.30E+13 | 150 | 9.75E+08 | Addgene | Th | 4 |
| ej3 | 466 | 7.2 | dual | PHP.eB | CAG | EGFP | 1.30E+13 | 100 | 6.50E+08 | Addgene | OB | 3 |
| ej4 | 466 | 8.9 | dual | PHP.eB | CAG | EGFP | 1.30E+13 | 100 | 6.50E+08 | Addgene | OB | 3 |
| ej5 | 466 | 9.1 | dual | PHP.eB | CAG | EGFP | 1.30E+13 | 100 | 6.50E+08 | Addgene | DP | 3 |
| ej7 | 176 | 6.2 | dual | PHP.eB | CAG | EGFP | 1.30E+13 | 200 | 1.30E+09 | Addgene | DP | 4 |
| ej8 | 209 | 7.5 | dual | PHP.eB | CAG | EGFP | 1.30E+13 | 200 | 1.30E+09 | Addgene | DP | 4 |
| ej13 | 657 | 10.9 | dual | PHP.eB | CAG | GCaMP8s | 2.06E+14 | 100 | 1.03E+10 | Neurotools | DP | 1 |
| ej14 | 657 | 11.4 | dual | PHP.eB | CAG | GCaMP8s | 2.06E+14 | 100 | 1.03E+10 | Neurotools | DP | 1 |
| ej15 | 701 | 8.9 | single | PHP.eB | CAG | GCaMP8s | 2.06E+14 | 100 | 1.03E+10 | Neurotools | DP | 1 |
| ej6 | 280 | 5.9 | single | PHP.eB | CAG | GCaMP8s | 2.10E+14 | 100 | 1.05E+10 | Neurotools | DP | 4 |
| ej9 | 329 | 10.2 | single | PHP.eB | CAG | GCaMP8s | 2.06E+14 | 100 | 1.03E+10 | Neurotools | DP | 0 |
| ej10 | 259 | 7.9 | single | PHP.eB | CAG | hM4D-mCherry | 4.49E+13 | 100 | 2.25E+09 | Neurotools | OB | 2 |
| ej11 | 259 | 8.2 | single | PHP.eB | CAG | hM4D-mCherry | 4.49E+13 | 100 | 2.25E+09 | Neurotools | OB | 2 |
| ej12 | 259 | 7.2 | single | PHP.eB | CAG | hM4D-mCherry | 4.49E+13 | 100 | 2.25E+09 | Neurotools | OB | 2 |
| ej18 | 144 | 4.9 | dual | PHP.eB | CAG | tdTomato | 2.40E+13 | 100 | 1.20E+09 | Addgene | DP | 4 |
| ej19 | 144 | 5.3 | dual | PHP.eB | CAG | tdTomato | 2.40E+13 | 100 | 1.20E+09 | Addgene | DP | 3 |
| ej21 | 194 | 7.8 | dual | PHP.eB | CAG | tdTomato | 2.40E+13 | 150 | 1.80E+09 | Addgene | Th | 0 |
| ej22 | 194 | 6.4 | dual | PHP.eB | CAG | tdTomato | 2.40E+13 | 150 | 1.80E+09 | Addgene | Th | 0 |
| ej25 | 319 | 7.74 | dual | PHP.eB | CAG | tdTomato | 2.40E+13 | 150 | 1.80E+09 | Addgene | DP | 4 |
| ej26 | 319 | 10.25 | dual | PHP.eB | CAG | tdTomato | 2.40E+13 | 150 | 1.80E+09 | Addgene | DP | 4 |
| ej13 | 657 | 10.9 | dual | PHP.eB | mDlx | NLS-mRuby2 | 4.70E+13 | 100 | 2.35E+09 | Neurotools | DP | 0 |
| ej14 | 657 | 11.4 | dual | PHP.eB | mDlx | NLS-mRuby2 | 4.70E+13 | 100 | 2.35E+09 | Neurotools | DP | 0 |
| ej16 | 710 | 9.4 | dual | PHP.eB | mDlx | NLS-mRuby2 | 4.70E+13 | 100 | 2.35E+09 | Neurotools | DP | 0 |

**Table S3.** Summary of injections in adult *Pleurodeles waltl* included in factor analysis of mixed data (FAMD). Continuous variables were age (days), weight (g), titer (v.g./mL), volume, total v.g., and expression score (0-4). Categorical variables were single vs. dual constructs injected, serotype, promoter, reporter, manufacturer and injection site (DP, dorsal pallium; OB, olfactory bulb; Th, thalamus).

| Pool name | Sequence |
|---|---|
| PHP-eB_GFP_B1_43PP | GAGGAGGGCAGCAAACGGAAACTTATCTACGTAGCCATGCTCTAG |
| PHP-eB_GFP_B1_43PP | TGTAGTTAATGATTAACCCGCCATGTAGAAGAGTCTTCCTTTACG |
| PHP-eB_GFP_B1_43PP | GAGGAGGGCAGCAAACGGAACTGAAACATAAAATGAATGCAATTG |
| PHP-eB_GFP_B1_43PP | AAAAACCTCCCACATCTCCCCCTGATAGAAGAGTCTTCCTTTACG |
| PHP-eB_GFP_B1_43PP | GAGGAGGGCAGCAAACGGAAATGGTTACAAATAAAGCAATAGCAT |
| PHP-eB_GFP_B1_43PP | TTGTTAACTTGTTTATTGCAGCTTATAGAAGAGTCTTCCTTTACG |
| PHP-eB_GFP_B1_43PP | GAGGAGGGCAGCAAACGGAAGTCCAAACTCATCAATGTATCTTAT |
| PHP-eB_GFP_B1_43PP | TTTTTCACTGCATTCTAGTTGTGGTTAGAAGAGTCTTCCTTTACG |
| PHP-eB_GFP_B1_43PP | GAGGAGGGCAGCAAACGGAAACGGTATCGATGCGGGGAGGCGGCC |
| PHP-eB_GFP_B1_43PP | GTCTGCTCGAAGCGGCCGCCCGGGTTAGAAGAGTCTTCCTTTACG |
| PHP-eB_GFP_B1_43PP | GAGGAGGGCAGCAAACGGAAAGGCGAAGACGCGGAAGAGGCCGCA |
| PHP-eB_GFP_B1_43PP | AGGGAGATCCGACTCGTCTGAGGGCTAGAAGAGTCTTCCTTTACG |
| PHP-eB_GFP_B1_43PP | GAGGAGGGCAGCAAACGGAACCGCTGGATTGAGGGCCGAAGGGAC |
| PHP-eB_GFP_B1_43PP | CCGGCAGCAGGCCGCGGGAAGGAAGTAGAAGAGTCTTCCTTTACG |
| PHP-eB_GFP_B1_43PP | GAGGAGGGCAGCAAACGGAAGGTGGCAACACAGGCGAGCAGCCAA |
| PHP-eB_GFP_B1_43PP | GCAGAAGGACGTCCCGCGCAGAATCTAGAAGAGTCTTCCTTTACG |
| PHP-eB_GFP_B1_43PP | GAGGAGGGCAGCAAACGGAAACGGAATTGTCAGTGCCCAACAGCC |
| PHP-eB_GFP_B1_43PP | AAGGACGATGATTTCCCCGACAACATAGAAGAGTCTTCCTTTACG |
| PHP-eB_GFP_B1_43PP | GAGGAGGGCAGCAAACGGAACGGCGATGAGTTCCGCCGTGGCAAT |
| PHP-eB_GFP_B1_43PP | CCCCTGTCCAGCAGCGGGCAAGGCATAGAAGAGTCTTCCTTTACG |
| PHP-eB_GFP_B1_43PP | GAGGAGGGCAGCAAACGGAAGAGCTGACAGGTGGTGGCAATGCCC |
| PHP-eB_GFP_B1_43PP | GAGGGGGAAAGCGAAAGTCCCGGAATAGAAGAGTCTTCCTTTACG |
| PHP-eB_GFP_B1_43PP | GAGGAGGGCAGCAAACGGAAGTGCACACCACGCCACGTTGCCTGA |
| PHP-eB_GFP_B1_43PP | CCAGTGGGGGTTGCGTCAGCAAACATAGAAGAGTCTTCCTTTACG |
| PHP-eB_GFP_B1_43PP | GAGGAGGGCAGCAAACGGAAGCAACCAGGATTTATACAAGGAGGA |
| PHP-eB_GFP_B1_43PP | CGGGCCACAACTCCTCATAAAGAGATAGAAGAGTCTTCCTTTACG |
| PHP-eB_GFP_B1_43PP | GAGGAGGGCAGCAAACGGAAATGATACAAAGGCATTAAAGCAGCG |
| PHP-eB_GFP_B1_43PP | AATGAAAGCCATACGGGAAGCAATATAGAAGAGTCTTCCTTTACG |
| PHP-eB_GFP_B1_43PP | GAGGAGGGCAGCAAACGGAATTAAGAATACCAGTCAATCTTTCAC |
| PHP-eB_GFP_B1_43PP | CCACATAGCGTAAAAGGAGCAACATTAGAAGAGTCTTCCTTTACG |
| PHP-eB_GFP_B1_43PP | GAGGAGGGCAGCAAACGGAATAAGCTTGATATCGAATTCTTACTT |
| PHP-eB_GFP_B1_43PP | TTTTGTAATCCAGAGGTTGATTATCTAGAAGAGTCTTCCTTTACG |
| PHP-eB_GFP_B1_43PP | GAGGAGGGCAGCAAACGGAAGGCGGCGGTCACGAACTCCAGCAGG |
| PHP-eB_GFP_B1_43PP | CAGCTCGTCCATGCCGAGAGTGATCTAGAAGAGTCTTCCTTTACG |
| PHP-eB_GFP_B1_43PP | GAGGAGGGCAGCAAACGGAATTGCTCAGGGCGGACTGGGTGCTCA |
| PHP-eB_GFP_B1_43PP | ATGTGATCGCGCTTCTCGTTGGGGTTAGAAGAGTCTTCCTTTACG |
| PHP-eB_GFP_B1_43PP | GAGGAGGGCAGCAAACGGAACGTCGCCGATGGGGGTGTTCTGCTG |
| PHP-eB_GFP_B1_43PP | AGTGGTTGTCGGGCAGCAGCACGGGTAGAAGAGTCTTCCTTTACG |
| PHP-eB_GFP_B1_43PP | GAGGAGGGCAGCAAACGGAACTCGATGTTGTGGCGGATCTTGAAG |
| PHP-eB_GFP_B1_43PP | GTGGTCGGCGAGCTGCACGCTGCCGTAGAAGAGTCTTCCTTTACG |
| PHP-eB_GFP_B1_43PP | GAGGAGGGCAGCAAACGGAAGCCATGATATAGACGTTGTGGCTGT |
| PHP-eB_GFP_B1_43PP | ACCTTGATGCCGTTCTTCTGCTTGTTAGAAGAGTCTTCCTTTACG |
| PHP-eB_GFP_B1_43PP | GAGGAGGGCAGCAAACGGAATGTTGCCGTCCTCCTTGAAGTCGAT |
| PHP-eB_GFP_B1_43PP | AGTTGTACTCCAGCTTGTGCCCCAGTAGAAGAGTCTTCCTTTACG |
| PHP-eB_GFP_B1_43PP | GAGGAGGGCAGCAAACGGAAGTCGCCCTCGAACTTCACCTCGGCG |
| PHP-eB_GFP_B1_43PP | CTTCAGCTCGATGCGGTTCACCAGGTAGAAGAGTCTTCCTTTACG |
| PHP-eB_GFP_B1_43PP | GAGGAGGGCAGCAAACGGAAAAGATGGTGCGCTCCTGGACGTAGC |
| PHP-eB_GFP_B1_43PP | GTCTTGTAGTTGCCGTCGTCCTTGATAGAAGAGTCTTCCTTTACG |
| PHP-eB_GFP_B1_43PP | GAGGAGGGCAGCAAACGGAAGCTGCTTCATGTGGTCGGGGTAGCG |
| PHP-eB_GFP_B1_43PP | CGGGCATGGCGGACTTGAAGAAGTCTAGAAGAGTCTTCCTTTACG |
| PHP-eB_GFP_B1_43PP | GAGGAGGGCAGCAAACGGAAGGTCACGAGGGTGGGCCAGGGCACG |
| PHP-eB_GFP_B1_43PP | GAAGCACTGCACGCCGTAGGTCAGGTAGAAGAGTCTTCCTTTACG |
| PHP-eB_GFP_B1_43PP | GAGGAGGGCAGCAAACGGAAAGGGTCAGCTTGCCGTAGGTGGCAT |
| PHP-eB_GFP_B1_43PP | AGCTTGCCGGTGGTGCAAATGAACTTAGAAGAGTCTTCCTTTACG |
| PHP-eB_GFP_B1_43PP | GAGGAGGGCAGCAAACGGAATGTGGCCGTTTACGTCGCCGTCCAG |
| PHP-eB_GFP_B1_43PP | CCTCGCCCTCGCCGGACACGCTGAATAGAAGAGTCTTCCTTTACG |
| PHP-eB_GFP_B1_43PP | GAGGAGGGCAGCAAACGGAACAGCTCCTCGCCCTTGCTCACCATG |

| | |
|---|---|
| PHP-eB_GFP_B1_43PP | GACCAGGATGGGCACCACCCCGGTGTAGAAGAGTCTTCCTTTACG |
| PHP-eB_GFP_B1_43PP | GAGGAGGGCAGCAAACGGAAGAGACAGCACAATAACCAGCACGTT |
| PHP-eB_GFP_B1_43PP | GCGGATCCAATTCTTTGCCAAAATGTAGAAGAGTCTTCCTTTACG |
| PHP-eB_GFP_B1_43PP | GAGGAGGGCAGCAAACGGAAATGAACATGGTTAGCAGAGGCTCTA |
| PHP-eB_GFP_B1_43PP | CAGGAGCTGTAGGAAAAAGAAGAAGTAGAAGAGTCTTCCTTTACG |
| PHP-eB_GFP_B1_43PP | GAGGAGGGCAGCAAACGGAACCCTCAAGGCTTTCACGCAGCCACA |
| PHP-eB_GFP_B1_43PP | CCCCCGCACAAAGGGCCCTCCCGGATAGAAGAGTCTTCCTTTACG |
| PHP-eB_GFP_B1_43PP | GAGGAGGGCAGCAAACGGAAGCTAATTACAGCCCGGAGGAGAAGG |
| PHP-eB_GFP_B1_43PP | AAGAAACAAGCCGTCATTAAACCAATAGAAGAGTCTTCCTTTACG |
| PHP-eB_GFP_B1_43PP | GAGGAGGGCAGCAAACGGAACGCGGTCAGTCAGAGCCGGGGCGGG |
| PHP-eB_GFP_B1_43PP | GTCCCGCCCGCTCACCTGTGGGAGTTAGAAGAGTCTTCCTTTACG |
| PHP-eB_GFP_B1_43PP | GAGGAGGGCAGCAAACGGAAGGGCGAAGGCAGCGCGCAGCGACTC |
| PHP-eB_GFP_B1_43PP | CGCGAGGCGGCGGCGGAGCGGGGCATAGAAGAGTCTTCCTTTACG |
| PHP-eB_GFP_B1_43PP | GAGGAGGGCAGCAAACGGAAGCCGCCGCCGCCGCCGCCTCGCCAT |
| PHP-eB_GFP_B1_43PP | CCCGCCGCGCGCTTCGCTTTTTATATAGAAGAGTCTTCCTTTACG |
| PHP-eB_GFP_B1_43PP | GAGGAGGGCAGCAAACGGAAATTGGCTGCCGCCGCACCTCTCCGC |
| PHP-eB_GFP_B1_43PP | AGGAAACTTTCGGAGCGCGCCGCTCTAGAAGAGTCTTCCTTTACG |
| PHP-eB_GFP_B1_43PP | GAGGAGGGCAGCAAACGGAACGTGGGGCTCACCTCGACCATGGTA |
| PHP-eB_GFP_B1_43PP | GGGGGAGATGGGGAGAGTGAAGCAGTAGAAGAGTCTTCCTTTACG |
| PHP-eB_GFP_B1_43PP | GAGGAGGGCAGCAAACGGAACAAGTAGGAAAGTCCCATAAGGTCA |
| PHP-eB_GFP_B1_43PP | GCGATGACTAATACGTAGATGTACTTAGAAGAGTCTTCCTTTACG |
| PHP-eB_GFP_B1_43PP | GAGGAGGGCAGCAAACGGAATACCGTCATTGACGTCAATAGGGGG |
| PHP-eB_GFP_B1_43PP | ACTGGGCATAATGCCAGGCGGGCCATAGAAGAGTCTTCCTTTACG |
| PHP-eB_GFP_B1_43PP | GAGGAGGGCAGCAAACGGAATGCCAAGTGGGCAGTTTACCGTAAA |
| PHP-eB_GFP_B1_43PP | ACTTGGCATATGATACACTTGATGTTAGAAGAGTCTTCCTTTACG |
| PHP-eB_GFP_B1_43PP | GAGGAGGGCAGCAAACGGAACCTATTGGCGTTACTATGGGAACAT |
| PHP-eB_GFP_B1_43PP | TCCACCCATTGACGTCAATGGAAAGTAGAAGAGTCTTCCTTTACG |
| PHP-eB_GFP_B1_43PP | GAGGAGGGCAGCAAACGGAACTAGAGCATGGCTACGTAGATAAGT |
| PHP-eB_GFP_B1_43PP | TCATTATTGACGTCAATGGTACTCTTAGAAGAGTCTTCCTTTACG |

**Table S4.** HCR probe sets
